# Segment Anything for Microscopy 2

**DOI:** 10.64898/2026.11.12.766082

**Authors:** Anwai Archit, Carolin Teuber, Titus Griebel, Lisa Marschhäuser, Beke Lohmann, Mostafa Aakhte, Nadine Vornberger, Sebastian von Haaren, Mette Handberg-Thorsager, Isabelle Bonne, Dominic Grün, Anne Beghin, Jan Huisken, Constantin Pape

## Abstract

Foundation models have advanced microscopy image segmentation, but existing models are trained on two-dimensional data, struggle with complex structures in volumetric data, and address only a narrow range of analysis tasks. Here we present Segment Anything for Microscopy 2 (μSAM2), a foundation model that adapts SAM2 to joint 2D and 3D segmentation in light and electron microscopy. Trained on annotations of more than 23 million cells and nuclei and 1.6 million cellular structures, μSAM2 achieves state-of-the-art automatic and interactive segmentation and outperforms generalist and specialist methods on volumetric data. Dedicated models further enable accurate mitochondrion and nucleus segmentation in electron microscopy. Beyond segmentation, μSAM2 supports pixel and object classification from few annotations as well as cell tracking in 2D and 3D time-lapse data. All functionality is available in the user-friendly μSAM napari plugin. We demonstrated the capabilities of μSAM2 on challenging use cases spanning developmental biology, organoid research, cardiac physiology, and histopathology.

## Main

Microscopy is a key technique for gaining insight into the processes that govern life across scales. Modern light microscopy (LM) can acquire large, high-resolution images of cells, tissues, organs, and whole organisms in fixed samples and over time^1–3^. High-content imaging applies LM at scale to cell cultures or tissue models to study the effects of interventions, such as genetic perturbations or compound treatments, under controlled conditions^4–6^. Volume electron microscopy (EM) can image cells and tissues at sub-organelle resolution^7–9^, enabling detailed analysis of cellular morphology and intercellular organization, up to the reconstruction of large neural circuits in EM connectomics^10–12^. The size and complexity of these data far exceed the capacity of manual analysis, creating a need for powerful, adaptable, and user-friendly image analysis solutions.

Image analysis can be divided into several tasks. Among the most important of which are instance segmentation (to find and delineate individual cells, nuclei, etc.), semantic segmentation (or pixel classification, to separate image pixels into different classes, such as into foreground vs. background or into different tissue types), object classification (to assign segmented objects to classes, e.g. for cell phenotyping), and cell tracking (to follow individual cells over time, including their divisions, to reconstruct cell lineages). Method development for these tasks began with the advent of digital imaging^13^. Early work established classical methods such as thresholding, convolutional filters, and morphological operations, which are routinely used through popular tools such as Fiji^14^. Classical image analysis can solve many simple tasks, such as foreground-background separation. However, it requires extensive per-image parameter tuning for more difficult tasks, such as cell instance segmentation, and fails to deliver satisfactory results for the most challenging ones, such as instance segmentation in 3D. These challenges have led to the adoption of machine learning, first through methods that train shallow classifiers on hand-crafted features^15^. Later, deep learning was adopted^16^, which can solve more difficult tasks and has enabled addressing large and challenging analysis problems^10,11,17–20^. However, this progress comes at the cost of higher computational demands and, more importantly in practice, the need for large amounts of labeled training data for the task at hand.

Creating such labeled data *de novo* for every analysis problem is not feasible in practice. Hence, generalist models have been introduced that are trained for a given task on large and diverse labeled datasets. For example, Cellpose^21^ and Mesmer^22^ for cell segmentation in LM, StarDist^23^ for nucleus segmentation in LM, HoverNet^24^ for nucleus segmentation in histopathology, and MitoNet^25^ for mitochondrion segmentation in EM. These tools can be directly applied to new data or further adapted by fine-tuning on a few labeled examples^26^. In addition, models trained for specific tasks are shared through a domain-specific model repository^27^ and can likewise be applied directly or fine-tuned.

More recently, foundation models for microscopy have emerged. They operate similarly to generalist models but address a wider range of modalities and/or analysis tasks. This follows the trend in general computer vision towards broadly capable models, which are trained either via self-supervised learning on massive datasets^28^, on paired image and text data^29,30^, or on large datasets with images and object labels^31,32^. Foundation models based on self-supervised and vision-language training have already been adopted in histopathology for clinical decision support^33–35^ and are now emerging for other modalities^36–39^. However, most microscopy foundation models outside of histopathology build on the Segment Anything Model (SAM)^31^ and adapt it to instance segmentation through further training on microscopy data with segmentation labels. These include Cellpose-SAM^40^ and CellSAM^41^ for cell and nucleus segmentation in LM, as well as our prior work μSAM^42^ for instance segmentation in LM and EM. Our PathoSAM^43^ applies the same approach to nucleus segmentation in histopathology. Compared to generalist models, foundation models yield higher-quality results across a wider range of imaging conditions. Furthermore, our models (μSAM and PathoSAM) support interactive segmentation, in which the model segments a target object, such as a cell, based on user input such as point or box prompts.

While foundation models yield highly accurate segmentation results in 2D, to the point that their performance has been claimed to be superhuman^40^, practical applications have shown that they struggle to segment structures with complex shapes or dense packing in volumetric data, where specifically trained networks remain superior^20,44,45^. This is likely because current foundation models were trained exclusively on 2D data and are applied to 3D data slice by slice, followed by postprocessing. Furthermore, current models address only a limited set of tasks: Cellpose-SAM and CellSAM support automatic instance segmentation, and μSAM supports automatic and interactive instance segmentation. However, many other analysis tasks are relevant in practice, such as segmenting different tissue types within an organ (semantic segmentation), determining the phenotypes of segmented cells (object classification), or tracking cells over time in live-cell microscopy. These tasks currently require dedicated models. Suitable training data or pretrained deep learning models are often lacking, so that large amounts of data must be annotated to obtain good results on difficult problems.

Here, we present μSAM2, a more capable foundation model for microscopy. It addresses these limitations through three major contributions:

1. A joint architecture and training procedure for 2D and 3D segmentation that adapts SAM2’s architecture^32^ to microscopy (**Fig. 1a**).
2. Three model families trained on large and diverse datasets: one for cell segmentation in LM and EM and nucleus segmentation in LM, one for mitochondrion segmentation in EM, and one for nucleus segmentation in EM (**Fig. 1b**).
3. Methods for automatic and interactive instance segmentation in 2D and 3D, for object and pixel classification based on our prior work^46^, and for cell tracking through integration of Trackastra^47^, a state-of-the-art tracking method (**Fig. 1c**).

**Figure 1:**
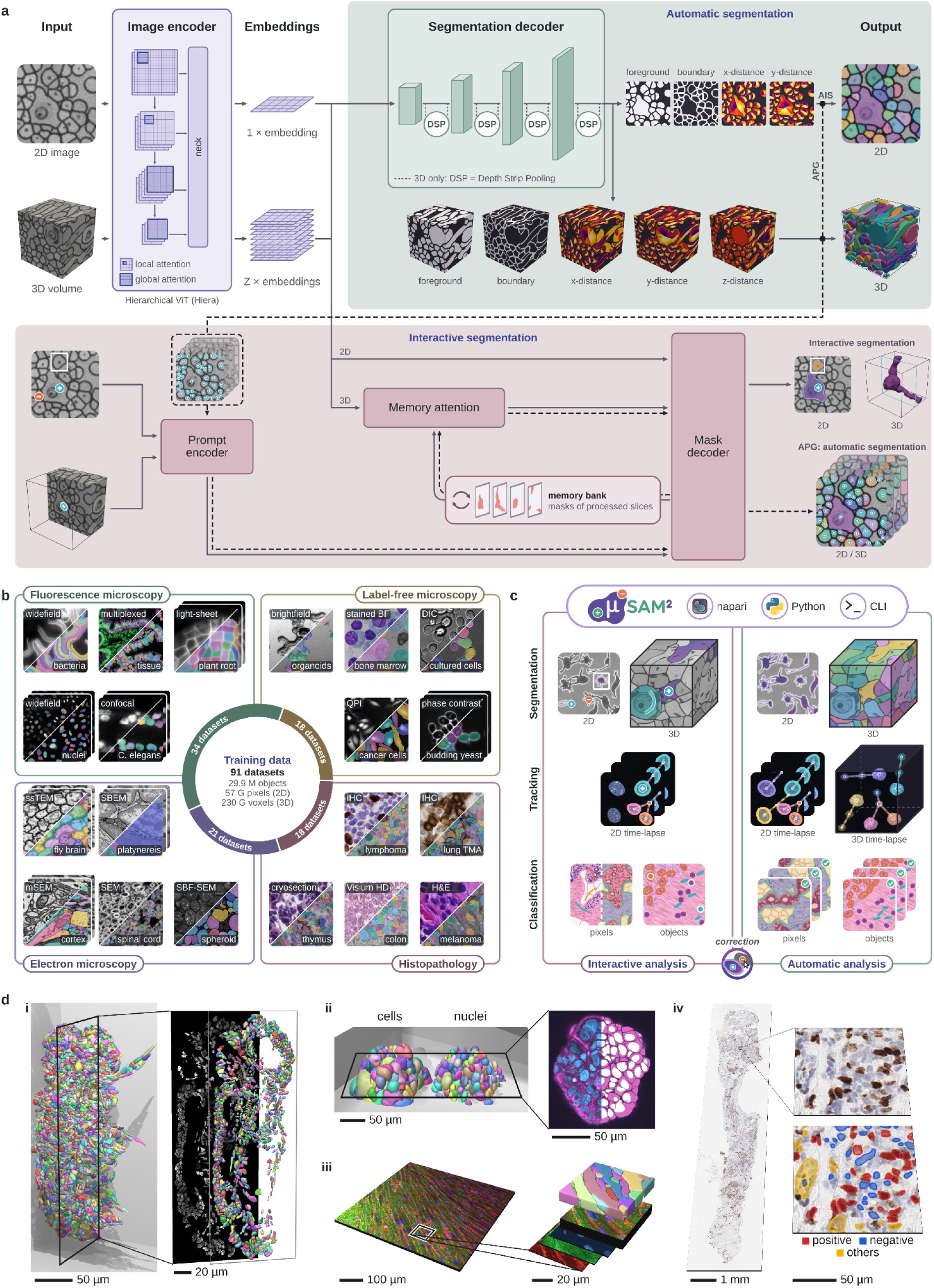
Contribution overview. **a**. μSAM2 architecture for 2D/3D interactive/automatic segmentation. **b**, training data of the μSAM2 models for cell, nucleus, and neurite segmentation. **c**, tasks addressed by the μSAM2 tool. **d**, overview of selected use-cases in this study: nucleus segmentation in an expanded *P. dumerilii* larva (i), cell and nucleus segmentation in an organoids (ii), cell segmentation and classification in cardiac tissue (iii), nucleus segmentation and classification in histopathology.

We integrated this functionality into our user-friendly μSAM tool so that it can be applied directly to diverse image analysis problems. We evaluated all contributions in rigorous benchmarks and found that our model is state-of-the-art for cell and nucleus segmentation in LM, outperforming prior generalist and foundation models on volumetric data. Furthermore, we found that our integrated functionality for classification and tracking yielded highly accurate results. Finally, we demonstrated how the tool can be applied to challenging analysis tasks in several use cases, ranging from developmental biology and organoid research to cardiac physiology and histopathology (**Fig. 1d**).

## Results

### Joint 2D and 3D training yields state-of-the-art microscopy segmentation

We designed a model architecture that can process both 2D and 3D input data. It is based on the hierarchical vision transformer of SAM2^32^ and retains SAM2’s components for interactive image and video segmentation. We extended it with a segmentation decoder that predicts a foreground mask and normalized geodesic distances for automatic segmentation (**Fig. 1a**, <u>Methods</u>). Next, we assembled a large and diverse training dataset from published microscopy data with segmentation labels for cells^20–22,26,48–76^, organoids^77–80^, and nuclei^48,49,74,81–94^ in LM, nuclei in histopathology^24,81,84,95–108^, and cellular structures (cells and neurites) in EM^10,11,17,109–124^ and modalities of similar appearance^19,125^. The dataset comprises both images and volumetric data. It contains annotations for 20,159,225 cells and 2,724,528 nuclei in 2D LM, 127,965 cells and 24,394 nuclei in 3D LM, and 1,618,484 cellular structures in EM (**Fig. 1b**, **Ext. Data Tab. 1 & 2**, <u>Methods</u>). We then implemented a training procedure that jointly trains this model for interactive and automatic segmentation on both the 2D and 3D data. We trained models of four sizes: *hvit_t_* (t for tiny), *hvit_s_* (s for small), *hvit_b_* (b for base), and *hvit_l_*(l for large) (<u>Methods</u>). Finally, we developed methods for interactive and automatic instance segmentation based on the trained models. For interactive segmentation, we use the model’s promptable segmentation functionality and repurpose its video segmentation mechanism for volumetric data. For automatic segmentation, we implemented two modes: *Automatic Instance Segmentation* (AIS), which uses the outputs of the segmentation decoder as input for a seeded watershed, and *Automatic Prompt Generation* (APG), which derives point prompts from the segmentation decoder outputs and passes them to the prompt encoder^126^. We integrated this functionality into the user-friendly μSAM tool, which is implemented as a napari plugin^127^ (<u>Methods</u>).

Next, we evaluated 2D automatic segmentation with μSAM2, stratified by cell segmentation in label-free microscopy, cell segmentation in fluorescence microscopy, nucleus segmentation in fluorescence microscopy, and nucleus segmentation in histopathology. For each modality, we used six *in-domain* (ID) datasets, corresponding to the test splits of datasets that are part of the training data of μSAM2 and of the most recent reference methods, and on six *out-of-domain* (OOD) datasets, which were not seen during training by any of the methods. This data comprises a total of 380,016 cells in ID^51,53,56,57,65,79^ and OOD^128–133^ label-free microscopy, 290,386 cells in ID^22,48,52,53,56,61^ and OOD^134–139^ fluorescence microscopy, 97,944 nuclei in ID^81,83–85,87,88^ and OOD^134,137,140–143^ fluorescence microscopy, and 121,909 nuclei in ID^95,97–99,105,106^ and OOD^144–149^ histopathology (**Ext. Data Tab. 1**). We compared to segmentation results from the general-purpose segmentation models SAM1–3^31,32,150^ and from the microscopy-specific models CellPose 3^151^, CellPoseSAM^40^, CellSAM^41^, CellViT++^152^, InstanSeg^153^, StarDist^23^, and μSAM^42^. We applied the domain-specific models only to the modalities they were designed for and used modality-specific variants where applicable (cell and nucleus model for CellPose 3, fluorescence and histopathology model for StarDist, fluorescence and histopathology^43^ model for μSAM). For SAM1 and 2, we used an approach that covers the image with a grid of points, uses each point as a prompt, and then assembles an instance segmentation via non-maximum suppression. For SAM3, we use the prompt *“cell”* for the concept-based segmentation mode of the model. All other models were applied with their default parameters. We computed the mean segmentation accuracy (mSA)^154^ (<u>Methods</u>). μSAM2 was the best method in six out of the eight different evaluation categories, the second best for ID cell segmentation in fluorescence microscopy (after CellPoseSAM), and the second best for OOD nucleus segmentation in fluorescence microscopy (after CellPose 3) (**Fig. 2 a**). Qualitative results showed accurate segmentation masks for diverse microscopy and histopathology image conditions (**Fig. 2 b**).

**Figure 2:**
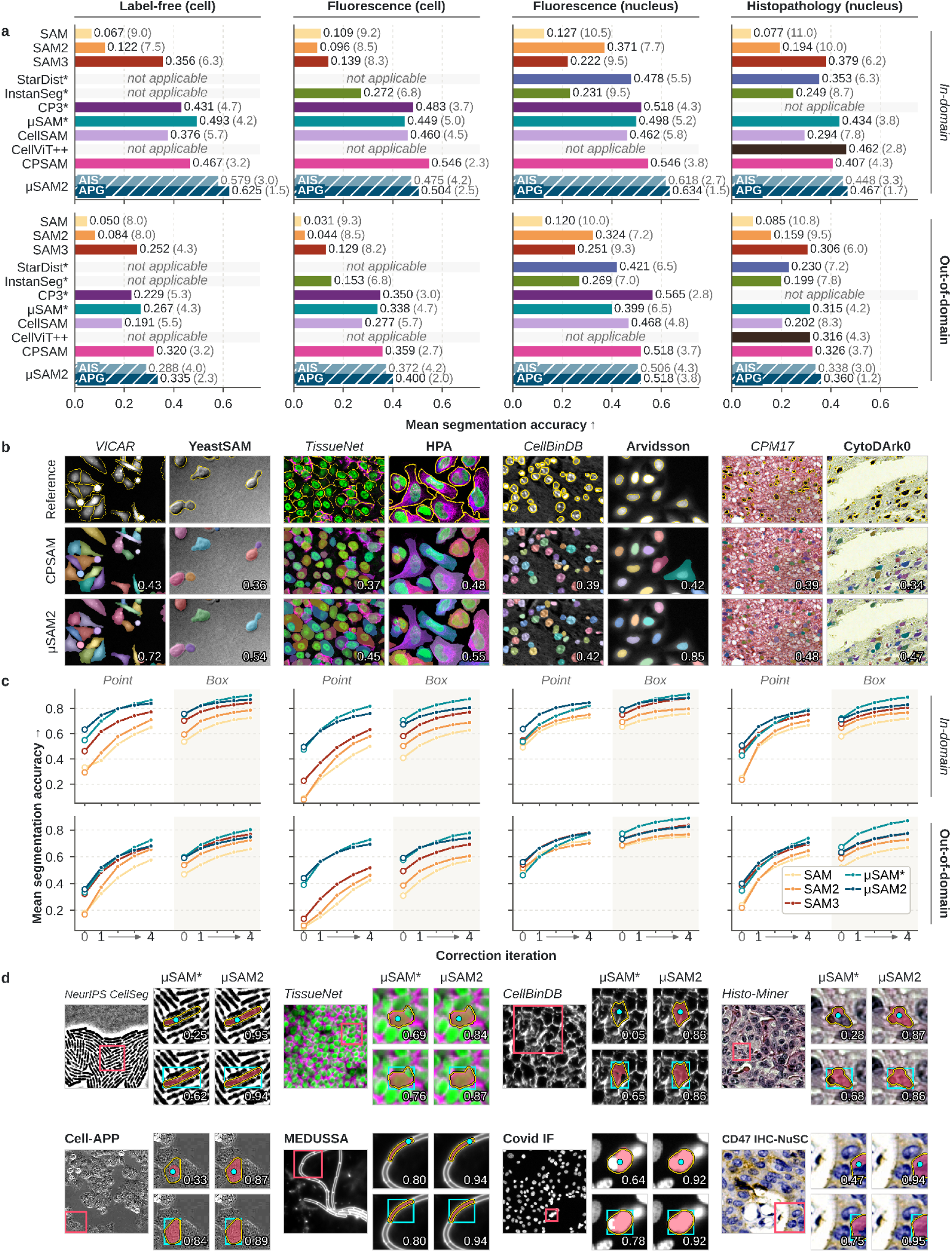
2D segmentation. **a.** Automatic segmentation results of μSAM2 and other methods, measured by mSA (higher is better); numbers in parentheses give the average rank, * marks methods using task-specific models. **b.** Example automatic segmentations of μSAM2 and CellPoseSAM; numbers give the per-image mSA. **c.** Interactive segmentation results of μSAM2 and other methods, starting from an initial point or box prompt (“0”) followed by four correction iterations (“1 → 4”). **d.** Example interactive segmentations of μSAM2 and μSAM, showing prompt (point / box), model output (red mask), and ground truth (yellow outline). Numbers indicate IoU between output and ground truth.

We then evaluated μSAM2 for interactive 2D segmentation, using the same evaluation data as before and compared to SAM1–3 and μSAM. For evaluation, we derived an initial point prompt or box prompt from a segmentation mask in the labels and then iteratively derived point prompts from errors. The masks were merged into an instance segmentation for the image and evaluated via mSA after each iteration (<u>Methods</u>). We found that μSAM2 performed better than the SAM models and on par with μSAM (**Fig. 2 c**). Visual inspection of the results showed that cell and nucleus shapes were captured well based on a single prompt. The masks from a single point prompt lead to partially segmented objects in some cases, while box prompts lead to more complete segmentations (**Fig. 2 d**).

Next, we evaluated automatic 3D segmentation. We compared to CellPose 3, CellPoseSAM, FOCUS-3D^44^, SegNeuron^109^, StarDist, and μSAM. We applied each model only to the tasks it was trained for. We used the cell- and nucleus-specific variants of FOCUS-3D and CellPose 3 and the default parameters for all models. For the CellPose models, we used overlap-based stitching of 2D segmentations, which we found to be superior to CellPose’s 3D mode that combines predictions along the three axes. We stratified the evaluation by ID^49,50,62,64,71,72^ and OOD^50,155,156^ data for cell segmentation in fluorescence microscopy (84,094 cells), ID^49,86,91,92,94^ and OOD^157–161^ data for nucleus segmentation in fluorescence microscopy (16,327 nuclei), and ID^11,19,111,116,124,162^ and OOD^10,12,109,163–165^ data for neurite segmentation in EM (93,978 neurites) (**Ext. Data Tab. 1**). μSAM2 outperformed the reference methods in all evaluations (**Fig. 3 a**). The Qualitative examples showed accurate μSAM2 segmentations across 3D imaging modalities (**Fig. 3 b**).

**Figure 3:**
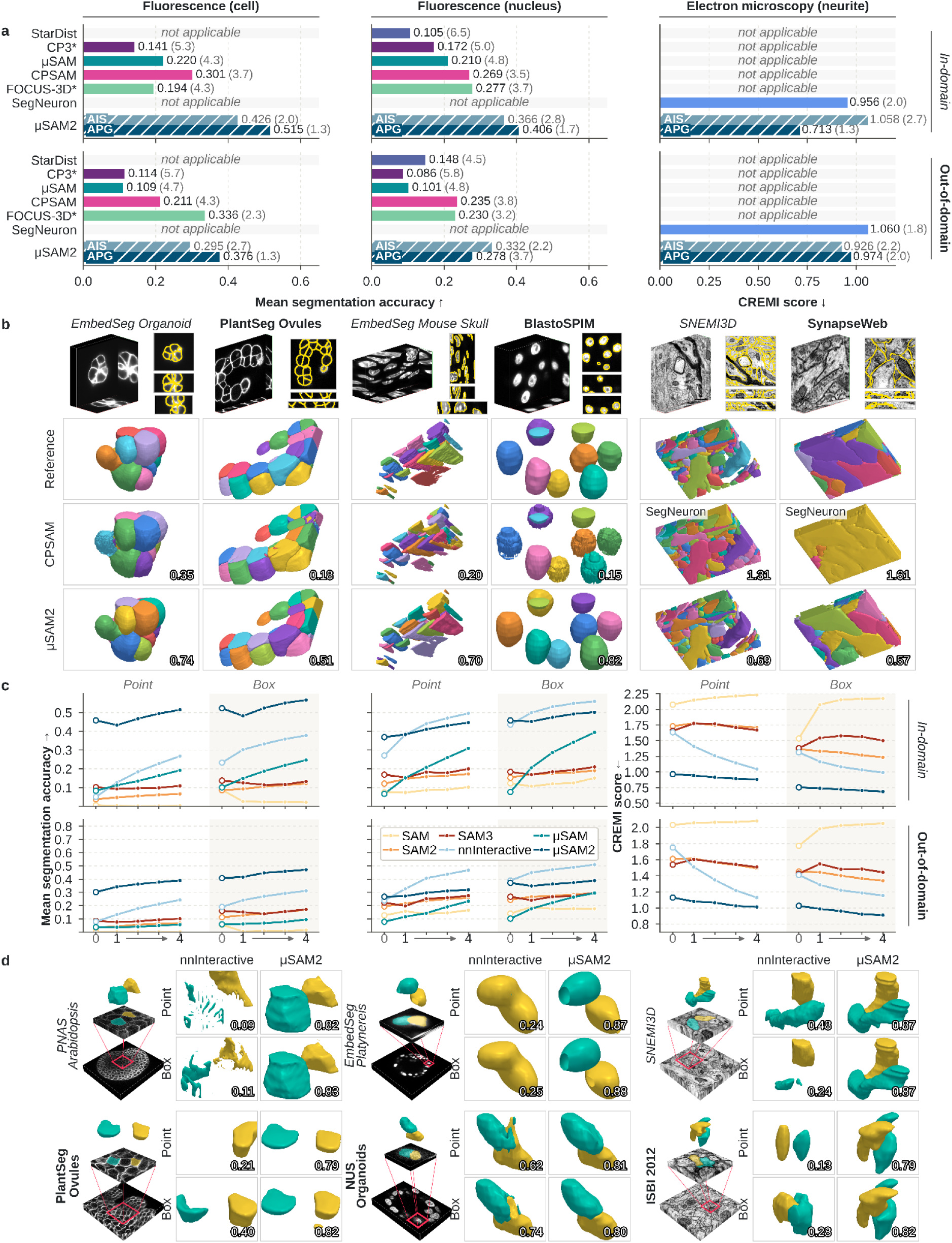
3D segmentation. **a.** Automatic segmentation results of μSAM2 and other methods, measured by mSA (higher is better); numbers in parentheses give the average rank and * marks methods using task-specific models. **b.** Example automatic segmentations of μSAM2, CellPoseSAM (for LM), and SegNeuron (for EM); numbers give the per-image mSA. **c.** Interactive segmentation results of μSAM2 and other methods, starting from an initial point or box prompt (“0”) followed by four correction iterations (“1 → 4”). **d.** Example interactive segmentations of μSAM2 and nnInteractive, showing two objects (cyan and yellow) in the ground truth and in the predictions of both models.

We also evaluated μSAM2 for 3D interactive segmentation on the same data. We compared to nnInteractive^166^, a model for interactive segmentation of volumetric biomedical image data, as well as to SAM1–3 and μSAM. We extended the 2D evaluation procedure to 3D: for each object, we derived an initial point or box prompt from its ground-truth mask in a single slice, segmented the object in 3D, and then iteratively added point prompts derived from the segmentation errors in the volume, evaluating mSA after each iteration (<u>Methods</u>). μSAM2 outperformed the other methods for cell segmentation in LM and neurite segmentation in EM. For nucleus segmentation in LM, μSAM2 performed best for the initial prompt but improved only modestly with correction prompts, so that nnInteractive performed better after correction. Both outperformed the remaining methods (**Fig. 3 c**). Qualitative examples show accurate 3D segmentations by μSAM2 from a single prompt (**Fig. 3 d**).

The results reported so far were obtained with the hvit_l_ model. We also evaluated the other model sizes, finding that segmentation quality improved with size as a general trend. The difference in automatic segmentation was more pronounced compared to interactive segmentation, where all models performed similarly (**Ext. Data Fig. 1**). Furthermore, we analyzed the runtimes for automatic and interactive segmentation, in 2D and 3D, on a CPU and GPU. For automatic segmentation in 2D, AIS was the fastest option, running in 0.9–1.8 seconds (s) per image on the CPU and 0.1s on the GPU, compared to 2.6s (CPU) and 0.2s (GPU) for CellposeSAM, and 4.0–4.8s (CPU) and 0.3s (GPU) for APG. We observed a similar trend for 3D, where AIS took 1.4–2.4s per slice on the CPU and 0.1s on the GPU, compared to 29s (CPU) and 1.5s (GPU) with CellposeSAM, and 70s (CPU) and 1.5s (GPU) for APG. For interactive segmentation, the image encoder took 0.3–1.3s on CPU and 0.01–0.04s on GPU (2D); 0.3s (GPU) and 0.01s per slice for 3D (**Ext. Data Fig. 2**). Application of the prompt encoder and mask decoder for per-object segmentation is considerably faster; the image encoder is applied only once per image, enabling interactivity.

Finally, we tested automatic 3D segmentation with μSAM2 on two challenging use-cases. First, we segmented the nuclei of a 6 day old *Platynereis dumerilii* larva that was expanded before imaging with a light-sheet microscope (**Fig. 1 d i,** <u>Methods</u>). We found the segmentation to be very accurate under visual inspection and measured 8,625 nuclei in the volume, which approximately followed a Gaussian distribution of nuclear volume (**Ext. Data Fig. 3**). Second, we segmented the cells and nuclei in organoids of breast tissue that were stained with a membrane and nuclear marker and imaged with confocal microscopy (**Fig. 1 d ii,** <u>Methods</u>). We found the segmentation to be accurate under visual inspection.

### μSAM2 embeddings enable interactive pixel and object classification

We extended μSAM2’s functionality to pixel classification (semantic segmentation) and object classification to support analysis tasks that are not addressed by instance segmentation, such as segmenting an organ into different tissue types or phenotyping already segmented cells. Here, we adopted the approach introduced in our prior work^46^. It uses the image embeddings predicted by μSAM2’s image encoder, or by another foundation model, as features for a random forest. Specifically, the embeddings are upsampled to the full image resolution and then either used as per-pixel features (pixel classification) or averaged over segmented object masks (object classification). This approach resembles ilastik’s pixel and object classification workflows15 but replaces the classical image features with embeddings from a foundation model. Similar approaches have previously been proposed for pixel classification^167,168^. We implemented this approach in new pixel and object classification widgets of the μSAM tool, where users can interactively provide labels, train a classifier, and inspect and correct its predictions (<u>Methods</u>).

We evaluated the object classification method for cell phenotyping on two histopathology datasets (PanNuke98: 5 classes, PUMA99: 10 classes) and on one hematology dataset (BBBC041129: 7 classes). We trained random forests based on object features derived from the foundation models DINOv3169, SAM1–3, μSAM, μSAM2 (hvit_t_), and UNIv233 (histopathology only), as well as on ilastik’s object features. The classifiers were trained on 25, 100, 1,000, or all objects sampled from the training split, applied to the respective test split, and evaluated using the mean F1-score across classes (<u>Methods</u>). We also trained a ResNet18170 as a supervised deep learning baseline. UNIv2 features outperformed all other methods on the histopathology datasets. For low label budgets, the other foundation models also outperformed ilastik features and the ResNet18. μSAM2 was among the best generally applicable models (excluding UNIv2) for all label budgets (**Fig. 4 a**). Qualitatively, classification based on μSAM2 features was reliable for common classes, yielding robust results when applied to larger tissue sections (**Fig. 4 c**).

**Figure 4:**
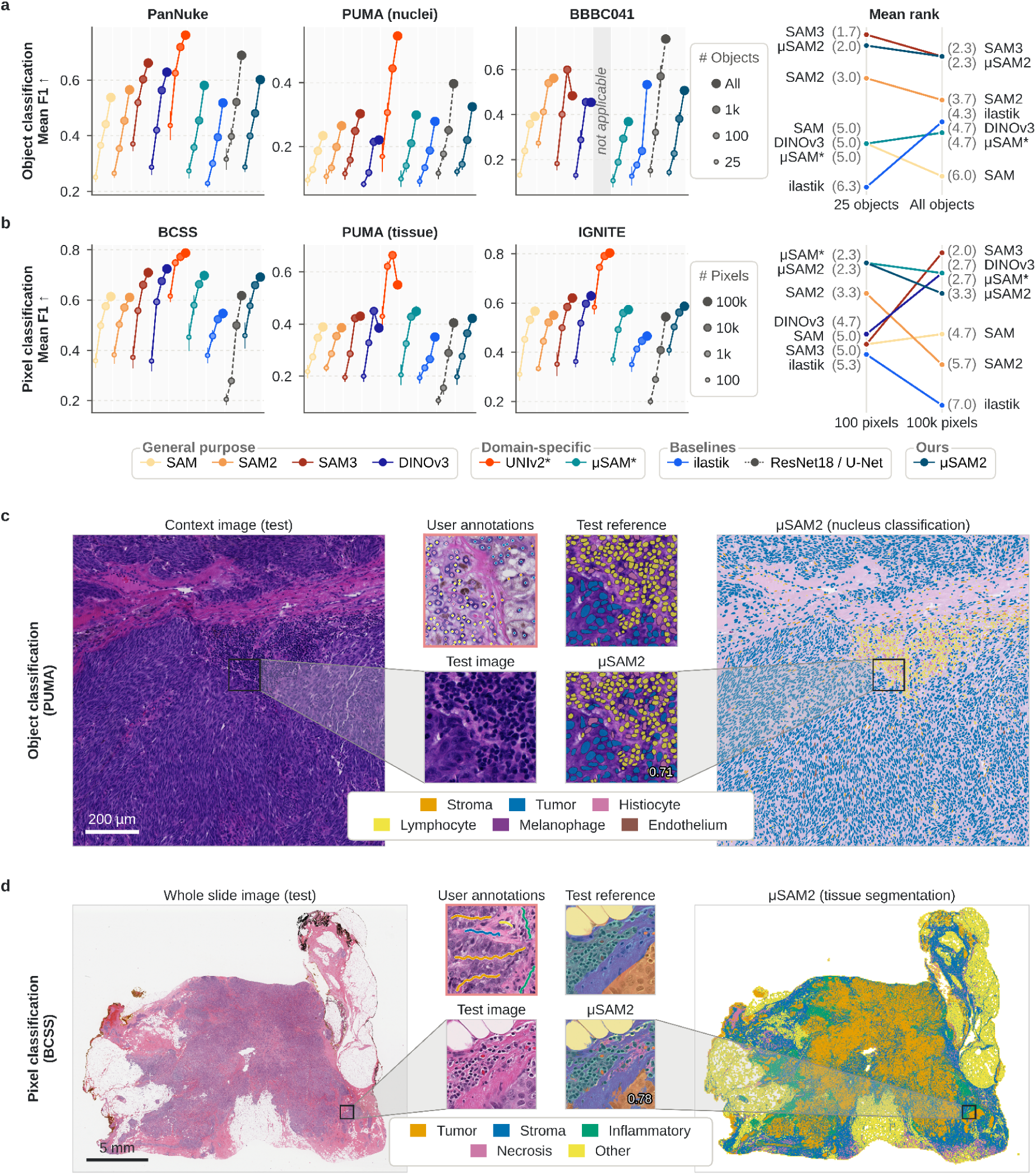
Object and pixel classification. **a**. Quantitative evaluation of object classification with a random forest trained using features derived from the embeddings of μSAM2 and other foundation models. Three datasets were used for evaluation under different annotation budgets (left), with mean ranks across datasets for the minimal and maximum budgets shown on the right. **b**. Quantitative evaluation of pixel classification, following a similar set-up as for object classification, using pixel-level features and brush stroke labels. **c.** Qualitative example of object classification with μSAM2 for nucleus phenotyping on the PUMA dataset^99^. **d**. Qualitative example of pixel classification with μSAM2 on the BCSS dataset^171^.

Next, we evaluated pixel classification on tissue segmentation in three histopathology datasets (BCSS171: 5 classes, PUMA^99^: 6 classes, IGNITE172: 6 classes). We followed a similar protocol as for object classification, deriving brushstroke labels from the ground-truth annotations and sampling training sets with 100, 1,000, 10,000, and 100,000 annotated pixels. We compared the same foundation models as before and used ilastik’s pixel features as well as a U-Net16 as baselines. UNIv2 again outperformed all other methods by a large margin, the other foundation models outperformed both ilastik features and the U-Net at all label budgets, and μSAM2 was again among the best generally applicable models across all label budgets (**Fig. 4 b**). Qualitatively, μSAM2 features yielded a robust whole-slide tissue segmentation from a modest annotation budget of 1,000 pixels (**Fig. 4 d**).

We then demonstrated the capability of the combined instance segmentation and object classification functionality of μSAM2 for two challenging use-cases. First, we turned to cardiac tissue imaged with confocal microscopy in 3D after staining with a membrane, ribosome, and nuclear marker (<u>Methods</u>). Here, we found the initial segmentations from μSAM2 to be of insufficient quality due to the complex, densely packed nature of cells in the volume. Hence, we annotated three small volumes (16 x 512 x 512 voxels) containing in total of 60 cells and then finetuned the hvit_l_ model on this data using the full training objective. This improved the segmentation quality substantially and made the results suitable for further analysis (**Fig. 1 d iii**). We will extend this use-case to 3D classification of cardiomyocytes and other cells in cardiac tissue.

Second, we analyzed 237 whole slide images (WSIs) of breast cancer tissue data from HISTAI173. The slides were immunohistochemically stained for the proliferation marker Ki67, which labels the nuclei of cycling cells brown via DAB, and counterstained with hematoxylin; we included only grade 3 tumors with a single reported Ki67 index (<u>Methods</u>). We segmented all nuclei with μSAM2 (APG), which yielded accurate results that required no further correction. We then extracted 60 crops and interactively classified the nuclei into Ki67-positive and Ki67-negative tumor cells and non-tumor cells using the object classification widget with μSAM2-derived features. This process took a total of 91 minutes and yielded annotations for 113,536 nuclei (1,737 user annotated) and an accurate classifier that we applied to all WSIs (**Fig. 1 d iv**). We then computed the local Ki67 index, i.e., the fraction of IHC-positive cells among the 500 nearest tumour cells, across regions with tumors. To focus on proliferation hotspots, we used the 90th percentile of local indices as the slide-level index. This index agreed well with the Ki67 index from the HISTAI pathology reports (intraclass correlation coefficient (ICC) 0.84, 95% CI 0.77–0.88; Spearman correlation 0.84, 95% CI 0.78–0.89). The ICC value on log-transformed value is 0.84 (95% CI 0.78–0.89), within the range that two studies report for inter-laboratory variability: 0.71 with local scoring methods^174^ to 0.87 with a standardized protocol^175^ (**Ext. Data Figure 4**).

### μSAM2 segmentation enables state-of-the-art cell tracking

We integrated Trackastra^47^, a state-of-the-art cell tracking model, with μSAM2 to enable the analysis of cell dynamics and development in live imaging. Trackastra tracks cells based on the image data and a given segmentation (here, from μSAM2). It supports tracking in both 2D and 3D time series. Automatic and interactive tracking, which relies on the promptable video segmentation module, are available in the tracking widget of the μSAM tool (<u>Methods</u>).

We evaluated automatic tracking with Trackastra, using segmentations from μSAM2 (AIS and APG), μSAM, and CellPoseSAM as input, on 2D time series (ID55,57,76,85 and OOD^131,139,142,176^) and 3D OOD time series^160^ with tracking annotations. We scored the results with the TRA, SEG, and LNK metrics^160^. TRA measures overall tracking accuracy (accounting for detection and linking errors), SEG measures segmentation quality (mean IoU of matched objects), and LNK measures linking accuracy alone. All metrics range from 0 to 1, with higher values being better (<u>Methods</u>). All approaches performed similarly on the 2D ID data, whereas μSAM2 with APG performed best on the 2D and 3D OOD data (**Fig. 5 a**). Qualitatively, μSAM2 produced fewer false links caused by segmentation errors than CellPoseSAM (**Fig. 5 b**).

**Figure 5:**
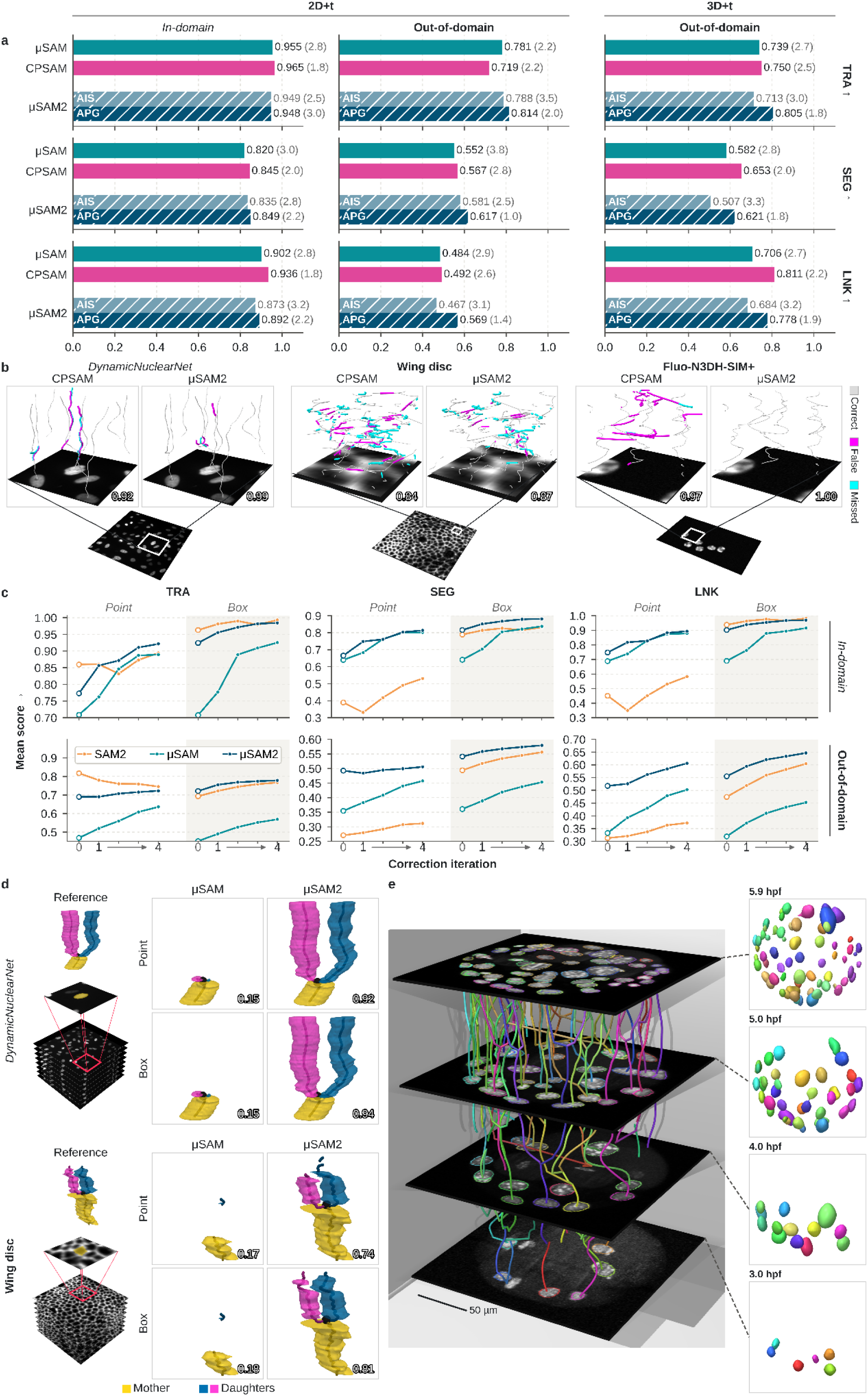
Cell and nucleus tracking. **a.** Automatic tracking with Trackastra based on segmentations from μSAM2 (AIS and APG), μSAM, and CellPoseSAM. Results are scored against reference annotations with TRA, SEG, and LNK^160^, which measure different aspects of tracking quality; higher is better for all. **b.** Examples of automatic tracking results based on μSAM2 (APG) and CellPoseSAM segmentations. Track colors indicate whether a track is correct or contains errors. **c.** Interactive tracking results for μSAM2, μSAM, and SAM2, evaluated with TRA, SEG, and LNK. Scores are averaged over tracklets (track segments between divisions) and datasets, for tracking from the initial point or box prompt (“0”) and after one to four correction iterations (“1 → 4”). **d.** Qualitative examples of interactive tracking of one lineage across a division. Each tracklet was obtained from a single prompt. **e.** Cell lineages of an early *P. dumerilii* embryo, reconstructed by automatic tracking with μSAM2 from a light-sheet microscopy time series.

Next, we evaluated interactive tracking on the same data, comparing μSAM2 with μSAM and SAM2. We split each lineage into tracklets (track segments between divisions), derived an initial point or box prompt from the annotation in the first frame of each tracklet, and then corrected the tracks in one to four iterations by adding point prompts derived from the annotations (<u>Methods</u>). μSAM2 performed best for most metrics; the exceptions were tracking accuracy (TRA) with initial point prompts on 2D data (ID and OOD) and with initial box prompts on 2D OOD data, where SAM2 performed better (**Fig. 5 c**). Qualitatively, tracklets predicted by μSAM2 from a single prompt appeared accurate (**Fig. 5 d**).

Finally, we applied μSAM2 to a challenging tracking problem: reconstructing cell lineages during the early development of a *P. dumerilii* embryo imaged with light-sheet microscopy (<u>Methods</u>). We segmented the nuclei in 3D with APG, which yielded results that appeared accurate on visual inspection. We then tracked these nuclei over time with Trackastra, which yielded plausible lineages (**Fig. 5 e**). We will extend this application to cell segmentation and to analyzing how cell morphology evolves across lineages.

### Dedicated models achieve state-of-the-art mitochondrion and nucleus segmentation in electron microscopy

We trained two additional μSAM2 models on EM data: one for mitochondrion segmentation and one for nucleus segmentation. We chose separate models because the nucleus training data contain unlabeled mitochondria and vice versa, which would yield inconsistent annotations in a joint training set. We assembled two training sets comprising 155,137 annotated mitochondria in EM images and volumes^10,25,119,162,177–194^, and 31,088 annotated nuclei in EM and X-ray images and volumes^10,17,120,186,189,195–204^ (**Ext. Data Tab. 1**, <u>Methods</u>). Using the same training procedure as before, we trained one hvit_s_ model per task (the other model sizes will be trained soon). These models support the same interactive and automatic segmentation functionality described above and are available in the segmentation and classification widgets of the μSAM tool.

We evaluated both models for 3D mitochondrion and nucleus segmentation on ID and OOD data comprising 14,484 mitochondria (ID^162,177–179,188,192^ and OOD^25,203–207^) and 3,127 nuclei (ID^10,17,186,189,197^ and OOD^114,193,205,208–210^). We compared to the EM model of μSAM, to MitoNet^25^, a method for mitochondrion segmentation in EM, and to NucleoNet^202^, a method for nucleus segmentation in EM. The input volumes for MitoNet and NucleoNet were rescaled to a voxel size that matched their respective training data’s voxel size. The evaluation followed the setup used for 3D LM data. The μSAM2 models outperformed all other methods by a wide margin, in particular in APG mode (**Fig. 6 a**). Visualization of exemplary results indicated a high segmentation quality for challenging 3D organelle segmentation tasks (**Fig. 6 b**).

**Figure 6:**
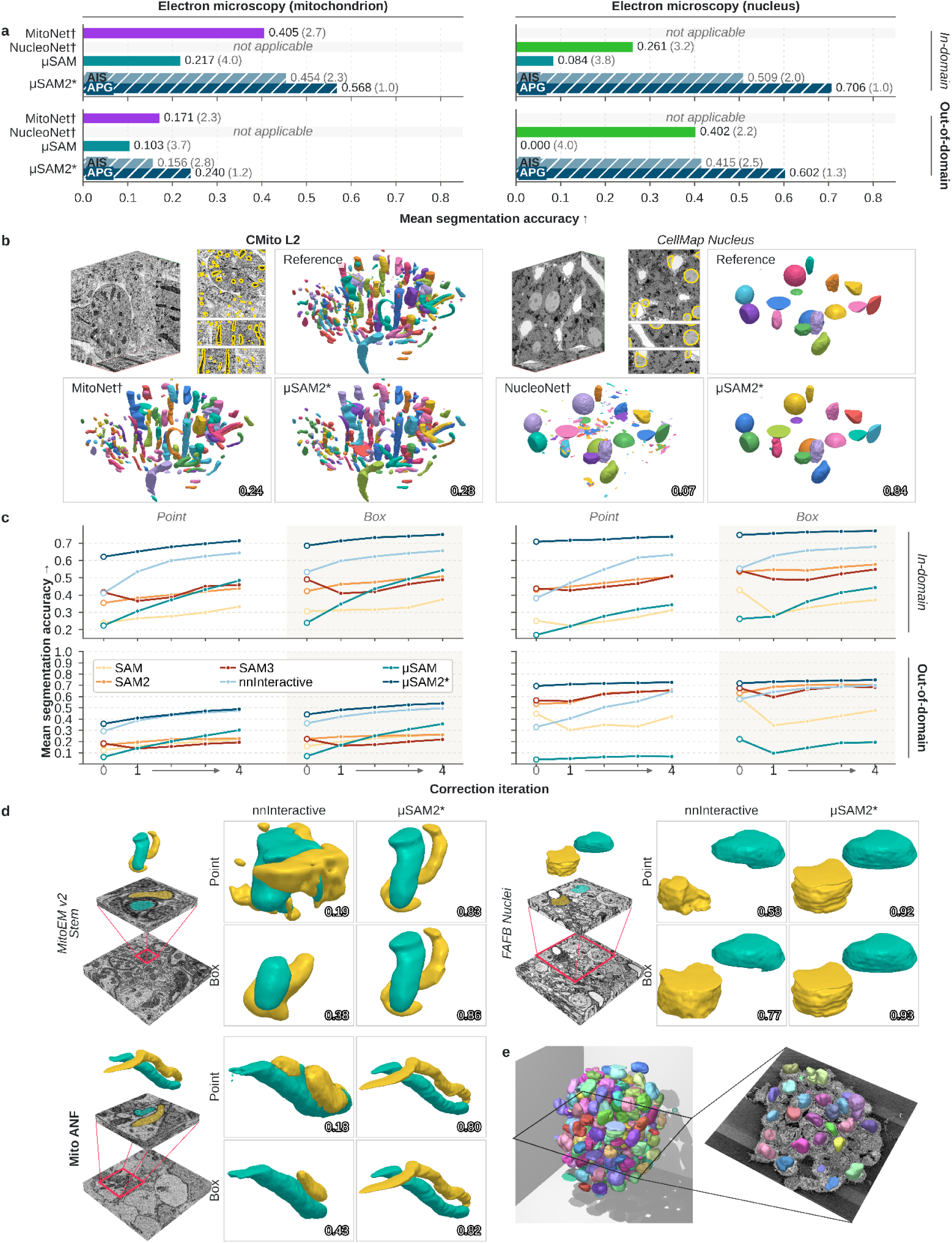
Organelle segmentation. **a.** Automatic segmentation evaluation of μSAM2 (mitochondrion and nucleus segmentation) compared to MitoNet, NucleoNet, and μSAM. Same evaluation set-up as Fig. 3 a. **b.** Qualitative examples for automatic segmentation results for mitochondrion and nucleus segmentation. **c.** Interactive segmentation evaluation, comparing μSAM2 to SAM1–3, μSAM, and nnInteractive. Same set-up as in Fig. 3 c. **d.** Qualitative examples of interactive segmentation results for organelles. **e.** Organoid imaged in volume EM with nucleus segmentation from μSAM2 (APG).

Next, we evaluated interactive segmentation on the same data, following the setup used for 3D LM data. We compared to nnInteractive, SAM1–3, and μSAM. The μSAM2 consistently performed better than the other methods (**Fig. 6 c**). Qualitative inspection showed accurate 3D segmentation results based on a single prompt (**Fig. 6 d)**.

Finally, we analyzed a colorectal organoid that was imaged in volume EM (<u>Methods</u>). We used the model for nucleus segmentation in APG mode to find and delineate the nuclei in the volume. We found that the nuclei were mostly segmented accurately. A few were missed in areas with low signal-to-noise ratio or imaging artifacts (**Fig. 6 e**).

## Discussion

We presented μSAM2, a foundation model for image analysis in microscopy that is available in a user-friendly tool. Its joint architecture and training procedure for 2D and 3D data distinguish it from prior work and lead to substantially better performance in 3D. We comprehensively evaluated automatic and interactive segmentation in 2D and 3D across a total of 28 different conditions. μSAM2 was the best method in the majority of these and it performed on par with μSAM in the eight conditions for 2D interactive segmentation. Another model performed better in only four conditions: CellPoseSAM for 2D automatic cell segmentation in fluorescence microscopy (ID), CellPose 3 for 2D automatic nucleus segmentation in fluorescence microscopy (OOD), and nnInteractive for 3D interactive nucleus segmentation in fluorescence microscopy (ID and OOD) after correction prompts.

Further, we integrated μSAM2 with Trackastra, resulting in a state-of-the-art approach for automatic cell tracking. In challenging evaluations, segmentations from μSAM2 yielded better tracking results than the other segmentations used as Trackastra input in 5 out of 9 tested settings. μSAM2 also supports interactive tracking, leveraging the video segmentation functionality of SAM2. Here, it performed better than the reference methods in 9 out of 12 tested settings.

For pixel and object classification, μSAM2’s features fell short of domain-specific foundation models, such as UNIv2 for histopathology. However, they are broadly applicable and were among the best features of the general-purpose foundation models we compared. The dedicated μSAM2 models for mitochondrion and nucleus segmentation in EM were the best methods in all eight settings we evaluated, covering automatic and interactive segmentation on ID and OOD data for both structures.

Finally, we applied μSAM2 to challenging use-cases across imaging modalities: we segmented 8,625 nuclei in an expanded *P. dumerilii* larva imaged with light-sheet microscopy, cells and nuclei in pancreatic organoids imaged with confocal microscopy, cells in cardiac tissue after fine-tuning on a small number of annotations, and nuclei in an organoid imaged with volume EM. We further derived a Ki67 index for 237 breast cancer WSIs that agreed well with pathologist scoring, based on a classifier that was trained in 91 minutes of interactive annotation, and reconstructed cell lineages in an early *P. dumerilii* embryo. These use-cases demonstrate that μSAM2 can solve diverse real-world analysis problems with modest user effort.

Overall, μSAM2 is the most broadly capable foundation model for microscopy image analysis. It will enable users to solve challenging analysis problems across domains much faster than currently possible. Its integration into the already popular μSAM tool ensures its distribution and promises broad adoption.

### Future developments

Several directions could further extend μSAM2. First, we already support fine-tuning, as shown in the cardiac tissue use-case, but have not yet studied and implemented efficient fine-tuning on user-grade hardware. Here, we can apply lessons from our prior work on parameter-efficient fine-tuning^211^, which showed that freezing all but the last layers of the image encoder provides the best trade-off between performance and efficiency. Second, our approach could be extended to other imaging modalities, in particular to medical imaging, building on our prior work212. We expect our training regime to improve on existing adaptations of SAM2 to medical images^213^. Third, a single pan-organelle model for EM that segments mitochondria, nuclei, the Golgi apparatus, the endoplasmic reticulum, and further organelles would be highly valuable. Such a model is currently not feasible because the required annotated training data is missing. A further promising direction is the integration with agentic systems that could leverage both μSAM2’s interactive and automatic segmentation functionality.

### Limitations

μSAM2 does not yet yield sufficiently accurate results for all supported segmentation tasks. In such cases, users can fine-tune the model. Annotating training data is fast thanks to the interactive segmentation functionality, but training requires powerful hardware and takes dozens of minutes to hours, even with parameter-efficient fine-tuning. This limits its use in a human-in-the-loop setting and generally makes fine-tuning more difficult. This limitation could be overcome with a few-shot learning approach, as has been demonstrated for instance segmentation with SAM3^150^.

Furthermore, users have to balance segmentation quality and runtime: APG is more accurate than AIS but slower, in particular on a CPU, and larger models generally have longer runtimes (**Ext. Data Fig. 2**).

Interactive cell tracking is not explicitly trained, even though μSAM2 inherits the corresponding training mechanism from SAM2 and suitable training data is available. Training for this task may further improve interactive tracking performance.

Finally, a fully fair comparison of segmentation methods is challenging due to differences in their training data, which means that ID and OOD data are not consistent across models. Here, we chose ID and OOD data to agree with most reference methods, but could not find ID data that was part of the training data of all other methods. Furthermore, how differences in the stringent evaluation metrics translate into biological relevance depends on the use-case; for many datasets, the results of several of the methods we compared to may already be adequate.

## Methods

The methods section gives an overview of μSAM2’s model architecture, its training data, and training procedure, followed by the implementation and evaluation of the instance segmentation, object and pixel classification, and tracking functionality. Finally, we explain the different use-cases that we included to demonstrate μSAM2’s capability. All of μSAM2’s functionality is available in the user-friendly μSAM tool, which is implemented as a napari plugin, and as python library and corresponding command line interface (CLI). The python library and CLI enable automatic segmentation, classification, and tracking via batch processing. The tool is available at https://github.com/computational-cell-analytics/micro-sam and well documented. The implementation is based on the PyTorch and SAM2 libraries, the torch_em, bioimage-cpp, and bioimage-py libraries we develop, and on libraries from the standard python scientific stack.

### Model architecture

μSAM2 is based on the model architecture of SAM2^32^ for prompt-based image and video segmentation. SAM2’s architecture consists of three main modules: the *image encoder*, the *prompt encoder*, and the *mask decoder*, complemented by memory modules for video segmentation. The image encoder is implemented as a hierarchical vision transformer (Hiera)^214^. It processes the input image, which is resized to 1024 × 1024 pixels, and outputs the *image embedding,* a 256-dimensional vector for each position of a 64 × 64 grid, corresponding to the image downsampled by a factor of 16. This embedding is computed only once per input. For video inputs, the image encoder is applied to each frame of the video independently. SAM2 is available in four variants with image encoders of different sizes: hvit_t_ (39M parameters), hvit_s_ (46M parameters), hvit_b_ (81M parameters), and hvit_l_ (224M parameters). The prompt encoder is implemented as a small artificial neural network that takes as input a combination of point coordinates, with either a positive (point belongs to the object of interest) or a negative (point does not belong to the object) label, bounding box coordinates, and masks. For video inputs, these prompts are distributed across frames. The prompt encoder maps the prompts to embedding vectors (prompt tokens). Image embedding and prompt tokens are inputs to the mask decoder, a small transformer-based neural network, which predicts the output mask for the object indicated by the prompts. This design enables efficient interactive segmentation because the image encoder, which contains most of the parameters, is applied only once per input image or video and its output can be reused for multiple objects and prompts. For video segmentation, SAM2 additionally contains a memory encoder, a memory bank, and a memory attention module (**Fig. 1 a**). The memory encoder fuses the image embedding of a frame with the mask predicted for it, and the memory bank stores the resulting memories of up to six previous frames as well as of the prompted frames. The memory attention module conditions the image embedding of the current frame on these memories before it is passed to the mask decoder. This temporal context enables tracking of objects along complex trajectories. SAM2 also supports the segmentation of multiple objects at a time in videos. The SAM2 models were trained on a large and diverse dataset containing 1.1 billion annotated objects in 11 million images and approximately 260 thousand manually annotated object trajectories in approximately 114 thousand videos.

We extended this architecture by introducing a decoder for automatic instance segmentation. It is implemented as a convolutional neural network that takes the image embedding as input and successively upsamples it to the resolution of the input image in four steps, each consisting of an interpolation that doubles the spatial resolution followed by convolutional layers. The image embedding is injected again through a skip connection after each upsampling step, following the design of μSAM’s segmentation decoder^42^. The same decoder processes 2D and 3D data: it is implemented with 3D convolutions, which reduce to 2D convolutions for 2D images by applying the central slice of each convolutional kernel. For 3D data, the image embeddings of the individual slices are stacked along the depth axis, so that the convolutions also exchange information between adjacent slices. In addition, each block is followed by depth strip pooling^215^, which averages the features along the depth axis and uses the result to rescale the features of each slice, thus spreading information across the whole volume. This operation is skipped for 2D data. The output block, which is shared for 2D and 3D data, consists of two convolutional layers followed by a 1×1 convolution. It predicts a channel with foreground probabilities, a channel with object boundary probabilities, and three channels with directed geodesic distances to the object center along the z, y, and x axes, of which the z channel is ignored for 2D data (**Fig. 1 a**). These channels serve as input for the instance segmentation operations (<u>Segmentation functionality</u>).

For interactive segmentation, we reuse SAM2’s prompt encoder, mask decoder, and memory modules. We process 3D data by recasting the depth axis as the time axis, which enables the bidirectional propagation of segmentations from prompted slices through the volume. Hence, μSAM2 supports interactive segmentation of 2D data, 3D data, and 2D time series data. We trained six model versions: four models with different encoder sizes for cell segmentation in LM and EM as well as nucleus segmentation in LM, one model with the hvit_s_ encoder for mitochondrion segmentation in EM, and one model with the hvit_s_ encoder for nucleus segmentation in EM. See the next two sections for a description of the <u>training data</u> and <u>training procedure</u>.

### Training data

#### Cell, neurite, and nucleus (LM) segmentation

We aggregated a large dataset from publicly available 2D and 3D microscopy data with instance annotations. Each dataset was inspected manually to ensure suitable data and annotation quality. For all datasets with a dedicated test split, we excluded this split from training; for several datasets without one, we held out a part of the data instead. We used these test data for ID evaluation (<u>Segmentation</u> <u>evaluation</u>). The dataset covers the following conditions (**Ext. Data Tab. 1 & 2**):

- Fluorescence microscopy: Cells and nuclei imaged in 2D and 3D, with widefield, confocal, two-photon, and light-sheet microscopy, as well as with multiplexed tissue imaging (e.g., CODEX, MIBI, Vectra). The data stem from diverse cell lines, tissues, and organisms, such as bacteria, yeast, epithelia, plant tissues, organoids, developing embryos and whole organisms (e.g., *Phallusia, P. dumerilii, C. elegans*), enteric neurons, as well as spiral ganglion neurons and hair cells in the cochlea.
- Label-free microscopy: Cells and organoids imaged in phase contrast, bright field, differential interference contrast (DIC) microscopy, as well as quantitative phase imaging (QIP). The data stem from diverse samples, such as bacteria, yeast, mammalian cell lines, blood smears, urine cytology specimens, and organoids derived from various organs.
- Histopathology: Nuclei in tissue sections stained with hematoxylin and eosin (H&E) or by immunohistochemistry (IHC), covering many human and mouse organs and cancer types. The images were extracted from whole-slide images, tissue microarrays, and cryosections.
- Electron microscopy: Neurites and cells imaged with volume EM (ssTEM, ssSEM, FIB-SEM, SBF-SEM, and multi-beam SEM) and electron tomography. The data stem from fly, mouse, human, and zebra finch brains, human platelets, tumor spheroids, and a *P. dumerilii* larva. For several large connectomics datasets, we used proofread automatic segmentations as annotations. We also included data from other techniques with an appearance and resolution similar to volume EM in this category: neurites imaged with expansion microscopy, and myelinated axons imaged with X-ray holographic nano-tomography and bright-field microscopy.

#### Mitochondrion segmentation (EM)

We aggregated a large dataset of mitochondria imaged in EM from 23 published datasets. The dataset contains 2D and 3D data acquired with different EM techniques: FIB-SEM, SBF-SEM, ssTEM, ssSEM, and 2D TEM. It covers tissues from fly, mouse, rat, and human, including brain (cortex, hippocampus, cerebellum, and brainstem), liver, skeletal muscle, urinary bladder, testis, and blood platelets, as well as cultured cells (e.g., HeLa, U2OS, and Jurkat cells) and colorectal cancer organoids. Some of these datasets only provide semantic labels, which we convert into instance labels using connected components or, for densely packed mitochondria, a seeded watershed. We also removed very small objects (fewer than 25 pixels in 2D or 50 voxels in 3D) from the labels. Otherwise, the data aggregation and processing procedure was the same as for the cell, neurite, and nucleus dataset.

#### Nucleus segmentation (EM)

We aggregated a large dataset of nuclei imaged in EM from 19 published datasets. The dataset contains predominantly 3D data acquired with different EM techniques, mainly FIB-SEM, ssTEM, SBF-SEM, array tomography SEM, and TEM. We also included data from soft X-ray tomography. The data stem from mouse tissues (e.g., pancreatic islets, liver, kidney, heart, and brain), the zebrafish heart, the fly brain, and a *P. dumerilii* larva, as well as from various cultured cells, spheroids, and organoids. Besides manual annotations, the dataset also contains proofread automatic segmentations, for example from large connectomics projects. The data aggregation and preprocessing procedure was the same as for the previous two datasets.

### Training procedure

We trained six models in total: four models of different sizes (hvit_t_ – hvit_l_) for cell, neurite, and nucleus segmentation in LM, and two models (hvit_s_) for mitochondrion and nucleus segmentation in EM. Each model was trained jointly for interactive and automatic segmentation in 2D and 3D. All components were initialized with the SAM2.1 weights, except for the segmentation decoder, which was randomly initialized. The interactive and automatic segmentation paths share the same image encoder.

In each training epoch, we drew a fixed number of samples from each dataset, with the number set per dataset according to its size. We extracted patches of 512 × 512 pixels at random positions from images larger than the patch size; smaller images were either padded or resized to the patch size, chosen at random. Patches containing fewer than three annotated objects (with a lower threshold for datasets with sparse objects) were rejected and redrawn. We normalized the input intensities using randomly sampled percentiles (lower percentile between 0 and 5, upper percentile set symmetrically) and applied random flips along all axes as further data augmentation.

Each batch contained either 2D or 3D data. For 2D iterations, we used batch sizes of 14 (hvit_t_), 12 (hvit_s_), 10 (hvit_b_), and 8 (hvit_l_). For 3D iterations, we used a batch size of 1, with volumes of 14 (hvit_t_, hvit_s_), 10 (hvit_b_), and 8 (hvit_l_) slices. The ratio of 2D to 3D iterations was set according to the ratio of 2D to 3D data in the respective training set. In each iteration, the interactive and automatic segmentation paths were trained one after the other, with separate backward passes.

The interactive segmentation path was trained following the procedure introduced by SAM2: Each image or volume slice was encoded independently by the image encoder, and a fixed number of objects (typically five) was sampled per iteration. For each object, an initial point or box prompt (each with 50% probability) was derived from its ground-truth mask, followed by seven correction rounds in which a positive and a negative point prompt were sampled from the ground truth to correct errors in the current prediction. This segmentation path was trained with a combination of Dice, focal, and binary cross-entropy losses applied to the predicted object masks. The additional output of the mask decoder that estimates mask quality was trained with a mean squared error loss. In 3D, the procedure was the same as in 2D, except that masks were propagated across the volume, using also the memory encoder and attention, and losses were computed for the volume.

The segmentation decoder of the automatic path predicts five channels: foreground probability, boundary probability, and the directed distances to the object center along x, y, and z. The corresponding targets (foreground mask, boundary mask, and normalized geodesic directed distances to the object center, defined as the maximum of the boundary distance transform) were derived from the ground-truth masks. The foreground and boundary channels were trained with a Dice loss and the distance channels with a mean squared error loss. The latter was applied only within the foreground mask. Training was the same in 2D and 3D, except that the distance along z was only trained in 3D.

We used the AdamW optimizer with an initial learning rate of 1e-5 and PyTorch defaults otherwise, together with the ReduceLROnPlateau learning rate scheduler conditioned on the validation loss (reduction factor of 0.9, patience of ten epochs). For validation, we drew up to 50 samples per dataset at the beginning of training and evaluated the same samples in every epoch, normalizing input intensities with the 2nd and 98th percentiles. The validation loss combined the Dice loss of the interactive path for a single prompt (point or box, drawn at random) with the full loss of the automatic segmentation path.

Each model was trained for approximately four days on eight H100 GPUs, using bf16 mixed precision and distributed training, corresponding to the following numbers of epochs:

- Cell, neurite, and nucleus models: 75 (hvit_t_), 73 (hvit_s_), 82 (hvit_b_), 79 (hvit_l_).
- EM: 140 (hvit_s_), for both the mitochondrion and nucleus model.

### Segmentation functionality

μSAM2 supports interactive and automatic segmentation of 2D images and 3D volumes. Interactive segmentation predicts object masks from user-provided prompts, whereas automatic segmentation derives a complete instance segmentation from the outputs of the segmentation decoder. Both build on the same image embeddings, which are computed once by the image encoder and shared between the two.

Interactive segmentation is implemented with μSAM2’s promptable 2D and 3D segmentation. It supports point prompts (positive and negative), bounding box prompts, mask prompts, which are drawn as polygons in the tool, and scribble prompts, which are converted into multiple point prompts (either positive or negative). Point and box prompts can be given for multiple objects at once, which are then segmented in a single batch; this is not supported for mask and scribble prompts or in combination with tiling. For 3D data, the masks are propagated through the volume in both directions; segmentation can also be restricted to a single slice or a selected range of slices.

For automatic segmentation, we implemented two modes, *Automatic Instance Segmentation* (AIS) and *Automatic Prompt Generation* (APG). Both operate as post-processing on the outputs of the segmentation decoder: the foreground probability map, the boundary probability map, and the normalized directed geodesic distances to the object center. Both modes support 2D images as well as 3D volumes.

For AIS, we implemented two variants. The first variant, *sparse,* is a seeded watershed. Its seeds are derived by tracing the flow defined by the directed geodesic distances to obtain a density map, which is then thresholded and labeled via connected components. The watershed uses the boundary predictions as heightmap and the thresholded foreground prediction as mask. This variant is well suited for images with background and for objects of limited spatial extent, such as cells or nuclei in light microscopy. In the second variant, *dense*, the watershed is followed by building a region adjacency graph over the watershed segments, deriving edge weights from the boundary channel predictions, and partitioning the graph into a segmentation via multicut^216^. This variant is well suited for densely packed, touching objects, such as neurites in volume EM, and can also segment large objects that extend throughout the image or volume.

The APG mode builds on our prior work^126^. It leverages promptable segmentation for automatic segmentation: it generates multiple candidate prompts per object, predicts the corresponding masks, and selects the best-fitting ones. The prompts are derived from the outputs of the segmentation decoder with the same procedure as the seeds in AIS, but with parameters adjusted such that multiple points are generated per object. Each point is passed as a positive prompt to the prompt encoder, and the mask decoder predicts the corresponding mask together with a quality estimate. Then, the masks are ranked by their quality estimate and merged into a consistent segmentation via non-maximum suppression. Finally, the masks can be derived by another set of prompts, such as a box derived from the first segmentation masks.

We selected the default parameters for the different segmentation modes based on a grid search that was run on the validation sets of the ID evaluation data.

The two modes offer different trade-offs between accuracy and runtime: AIS is fast and yields reliable results for many tasks, whereas APG is more accurate and thus particularly suited for difficult segmentation tasks. However, APG is slower: its runtime scales with the number of prompts and is especially long on CPU (<u>Results</u>).

Interactive and automatic segmentation for 2D and 3D data are integrated in a single widget, the *Segmentation Annotator*, in the napari plugin. The embeddings can be precomputed and saved, so that annotation can be resumed later without recomputing them. The *Batch Annotator* widget enables processing of multiple images or volumes sequentially.

The tool supports image data with a single channel or with three channels, which must be ordered as a channel with nuclear stain, membrane stain, and an optional further stain (set to zero if not given), or histopathology stained data with the usual RGB channel orders. All operations support tiling with overlap to process large images and volumes, with a default tile size of 512 × 512 pixels as the inner tile, with an additional overlap of 128 x 128 pixels; the tiling parameters can be adjusted. For automatic segmentation, prediction and prompt generation (for APG) are parallelized over tiles, followed by post-processing across tiles to obtain a consistent segmentation. For interactive segmentation, the mask is predicted in the tile containing the prompt and, if needed, propagated to adjacent tiles. The hyperparameters for all operations are set to default values (see above) and can be adapted by the user in the advanced settings.

### Segmentation evaluation

We evaluated segmentation results with the mean segmentation accuracy (mSA)^154^ for cells, nuclei, and mitochondria. To compute it, predicted and ground-truth objects are matched based on their intersection over union (IoU) at thresholds t from 0.5 to 0.95 in steps of 0.05. For each threshold, the segmentation accuracy is computed as SA(t) = TP(t) / (TP(t) + FP(t) + FN(t)), where TP is the number of matched objects (true positives), FP the number of unmatched predicted objects (false positives), and FN the number of unmatched ground-truth objects (false negatives). The mSA is the mean of SA(t) over all thresholds. It ranges from 0 to 1, with higher values indicating better results. For neurite segmentation in EM, in 2D and 3D, we instead used the CREMI score^124^. It is the geometric mean of the variation of information and the adapted Rand error. It is 0 for a perfect segmentation, with lower values indicating better results. This measure is more appropriate for dense segmentation tasks with many, complex-shaped objects, such as long neurites, where object-matching based measures like the mSA are too punitive.

We evaluated segmentation on a total of 93 datasets, which we grouped into nine categories according to dimensionality, imaging modality, and segmented structure. For each category, we used in-domain (ID) datasets, corresponding to the test splits of datasets that are part of the training data of μSAM2 and of the most recent reference methods, and out-of-domain (OOD) datasets, which were not used for training by any of the methods (**Ext. Data Tab. 1 & 2**). All evaluations were performed on 2D images of 512 × 512 pixels and on 3D volumes of 32 × 512 × 512 voxels, which we extracted from different images or volumes across the test data of each dataset.

For automatic segmentation, we compared μSAM2 to general-purpose and microscopy-specific segmentation models. For μSAM2, we evaluated both automatic segmentation modes, AIS and APG. As general-purpose models, we used SAM and SAM2, in both cases the large model variant with automatic mask generation, which covers the image with a regular grid of point prompts and assembles the predicted masks into an instance segmentation via non-maximum suppression, as well as SAM3, with the text prompt “cell” in its concept-based segmentation mode. As microscopy-specific models, we used CellPose 3 (cyto3 and nucleus model), CellPoseSAM, CellSAM, CellViT++, InstanSeg, StarDist (fluorescence and histopathology models), μSAM (vit_l_ models for light microscopy and histopathology), FOCUS-3D (cell and nucleus models), and SegNeuron for neurite segmentation in EM. We applied the microscopy-specific models only to the modalities and structures they were trained for. For 3D segmentation with CellPose 3 and CellPoseSAM, we segmented each slice in 2D and stitched the results based on overlap, which we found to be superior to CellPose’s 3D mode that combines predictions along the three axes. For EM organelle segmentation we used MitoNet, NucleoNet, and the μSAM organelle model (vit_l_ as reference methods). For all models, we used the default parameters and only adapted the minimum object size to each evaluation dataset.

For interactive segmentation, we simulated user annotations by deriving prompts from the ground-truth labels. Each object was prompted individually. The initial prompt was either a single positive point or a bounding box. The bounding box was fit tightly around the object; the point was placed at the eccentricity center of the object, which lies inside the object also for irregular shapes. We then performed four correction iterations, each adding one positive and one negative point prompt derived from the errors of the previous prediction with respect to the ground truth. The positive point was sampled at random in the missing region of the object or, if there was none, in the correctly segmented region. The negative point was sampled at random in the false positive region or, if there was none, in the region outside the dilated object mask. For the SAM2-based models, we additionally passed the mask logits of the previous prediction as a prompt in each correction iteration. We used the same random seed for all methods, so that the same initial prompts were sampled. For 3D data, the initial prompt was placed in the central slice of the object along the z-axis and the mask was propagated through the volume in both directions; the correction prompts were placed in the slice with the most errors, after which the object was re-propagated through the volume. After the initial prompt and after each correction iteration, the object masks were merged into an instance segmentation of the image or volume and evaluated with the measures described above. We compared to SAM1–3 and μSAM in 2D, and additionally to nnInteractive in 3D.

### Classification functionality

We implemented interactive object and pixel classification based on our prior work^46^. For object classification, we start from a given instance segmentation (from μSAM2 or another source) and image embeddings. Per-object features are computed by upsampling the embedding vector to the image size, averaging each embedding dimension across the object mask, and appending the object size as an additional feature. Labels for semantic classes are derived from user brushstrokes that are mapped to the instances. A random forest is trained based on these features and instances and applied to all instances in the image. Training and prediction take only a few seconds on a CPU and thus support interactive annotation and correction.

Pixel classification operates similarly. Here, embeddings are upsampled to the image size using AnyUp^217^ and the embedding vector for each pixel is used as a feature. User brushstrokes provide semantic labels on the pixel level, followed by random forest training and prediction of the full image or volume.

We implemented the *Object Classifier* and *Pixel Classifier* widgets in the napari plugin to provide the functionality to users. Both work on 2D and 3D data. They use μSAM2 as the default model and also support features derived from the following other vision foundation models: SAM1–3, μSAM models, MedicoSAM^212^, DINOv3^169^, and UNIv2^33^. These widgets can also be run in a batch mode, where they are applied to a series of images and volumes. The classifier state is kept, so that a larger training set can be built up. The classifier can be saved and then be applied to other data automatically using the CLI or python library.

### Classification evaluation

We evaluated the classification functionality using the protocol introduced in our prior work^211^. For object classification, a training set with either 25, 100, 1k, or all objects is sampled from a pool of training images, using class-balanced sampling. These annotations are used to train the classifier, which is then applied to the test set. The results are evaluated with the F1-score averaged over classes. We used the PanNuke^98^ (5 classes, histopathology), PUMA^99^ (10 classes), and BBC041^129^ (7 classes, hematology) datasets for evaluation and compared to features derived from SAM1–3, μSAM, DINOv3, and UNIv2 (histopathology only) embeddings. In addition, we compared to a ResNet18^170^ with ImageNet pretrained weights trained in a supervised fashion and ilastik^15^ object classification features.

We used a similar set-up for pixel classification, using 100, 1k, 10k, 100k labeled pixels. We evaluated on the three histopathology datasets BCSS^171^ (5 classes), PUMA (6 classes), and IGNITE^172^ (6 classes). We used the same vision foundation models for comparison as before and used a U-Net^16^ trained via supervised learning and ilastik pixel classification features as further references.

### Tracking functionality

The automated tracking functionality of μSAM2 uses Trackastra^47^, a transformer-based cell tracking method. Briefly, Trackastra takes a time series of images or volumes and the corresponding segmentation as input, computes simple morphological and intensity features for each segmented object, and feeds them into a transformer that predicts association scores between objects in nearby frames. These scores define the costs of a linking problem, which is solved either with a greedy heuristic or as an integer linear program. We integrated Trackastra into μSAM (in the Python library, the command-line interface, and the tracking widget of the napari plugin), using the segmentation obtained with μSAM2 as its input. Trackastra provides multiple pretrained models.

The tracking widget also supports interactive tracking of 2D data, which builds on the video segmentation functionality that μSAM2 models inherit from SAM2. Users prompt a cell in the frame in which it first appears, the model segments the corresponding tracklet by propagating the segmentation forward in time, and the result can be corrected interactively with additional prompts. The model does not detect cell divisions; instead, the user marks each division with a dedicated indicator, after which the daughter cells are tracked as separate tracklets.

Automatic and interactive tracking can be combined: users can first track all cells automatically, then remove incorrect tracks and re-track the affected cells interactively. Tracking results can be exported in the GEFF^218^ and CTC^160^ file formats.

### Tracking evaluation

We evaluated tracking results with the TRA, SEG, and LNK scores, which measure overall tracking quality, segmentation quality, and linking quality, respectively. All three measures were defined by the Cell Tracking Challenge^160^. TRA is based on the acyclic oriented graph matching (AOGM) measure, i.e., the weighted number of operations needed to transform the predicted tracking graph into the ground-truth graph (adding, removing, or splitting objects, and adding, removing, or relabeling links). It is normalized as TRA = 1 − min(AOGM, AOGM₀)/AOGM₀, where AOGM₀ is the cost of creating the ground-truth graph from scratch, and ranges from 0 to 1, with higher values indicating better results. LNK is computed in the same way but only considers linking errors, so it measures linking quality independently of detection errors. SEG is the mean Jaccard index between each ground-truth object and its matching predicted object; a predicted object matches a ground-truth object if it covers more than half of the ground-truth object’s pixels, and unmatched ground-truth objects contribute a score of 0. We computed TRA and LNK with the traccuracy library, available at https://github.com/live-image-tracking-tools/traccuracy.

To evaluate automatic tracking, we used segmentations from μSAM2, μSAM, and Cellpose-SAM as input to Trackastra and compared the resulting tracks. We used the *general_2d_w_SAM2_features* model for 2D+t data and the *ctc* model for 3D+t data, in both cases with the greedy linking solver. The following datasets were used for evaluation (**Ext. Data Tab. 2**):

- 2D+t:

○ 4 ID datasets: TOIAM, YeaZ, Cell-ACDC, DynamicNuclearNet^55,57,76,85^
○ 4 OOD datasets: ALFI, Cardioblast, EColi, WingDisc^131,139,142,176^
- 3D+t:

○ 6 OOD datasets from the Cell Tracking Challenge: Fluo-C3DH-A549-SIM, Fluo-C3DH-H157, Fluo-C3DL-MDA231, Fluo-N3DH-CHO, Fluo-N3DH-SIM+, Fluo-N3DL-TRIC

For interactive tracking, we compared μSAM2, μSAM, and SAM2 on the same 2D+t datasets (interactive tracking is not supported for 3D+t data). We split each lineage into tracklets and segmented each tracklet with prompts derived from its annotation: we started with a box or point prompt in the first frame of the tracklet and then performed four correction iterations, each adding a positive and a negative point prompt derived from the segmentation errors. The combined result for all tracklets of a time series was then scored with the metrics described above.

### Use-cases

Here, we describe the data and methodology for the different use-cases that we have included to demonstrate μSAM2’s capability on challenging, real world examples.

#### Platynereis larva (expansion, light-sheet microscopy)

Tissue expansion was performed on the marine larva *Platynereis dumerilii* (Annelida) following the protocol for ChromExM^219^. A 6-day-old larva was directly fixed for 15 min at room temperature, with shaking, in a fixative (3% paraformaldehyde (PFA, E15710, Electron Microscopy Sciences), 0.1% glutaraldehyde (16019, Electron Microscopy Sciences) in PBS), followed by 3 washes in PBS. The larva was post-fixed in 1% acrylamide and 0.7% PFA at room temperature for 6 hours, with shaking, and then washed briefly 2x in PBS. The larva was embedded in a gelling solution and expanded in two rounds. The first gelation and expansion step led to a ∼3.5x expansion. The second gelation and expansion step led to a total of ∼10x tissue expansion. The nuclei were counterstained with SYTOX Green (1:5000, S7020, Thermo Fisher) in Hanks’ Balanced Salt Solution (HBSS) (14025092, Thermo Fisher) after the final expansion. Following expansion, the hydrogel containing the larva was trimmed into a cube of approximately 5 × 5 × 5 mm and secured in a custom 3D-printed clamp. The sample was imaged in Milli-Q water using a custom-built isotropic light-sheet microscope^1^ configured for water immersion. The microscope was equipped with an air excitation objective (20×, NA 0.4; Mitutoyo), a multi-immersion detection objective (54-10-12, nominal NA 0.4; Applied Scientific Instrumentation) and a pco.panda 4.2 sCMOS camera (Excelitas PCO). The expanded sample was excited with a 488 nm laser line, and images were acquired with an exposure time of 25 ms. The entire specimen was acquired as 10 tiles arranged in 2 columns × 5 rows with ∼25% overlap. The voxel size was isotropic at 0.42 µm, with the axial step matched to the lateral image pixel size and the light-sheet thickness. We then stitched the tiles in BigStitcher^220^, using phase correlation to compute pairwise shifts followed by global optimization with rigid transformations. For faster handling, the tiles were fused into a single volume downsampled 3-fold (1.26 µm voxels). All scale bars in the figures were rescaled by the expansion factor and therefore show pre-expansion (biological) dimensions.

#### Organoids (confocal microscopy)

Organoids were grown from Mcf7 cells, a breast cancer cell line. The experimental protocol followed Ong et al.^161^: Cells were seeded into Lipidure-passivated JeWell microwell arrays in 24-well plates at about 20–50 cells per microwell. They were cultured for one week in DMEM high glucose supplemented with 10% FBS and penicillin–streptomycin, with the medium changed every three days. The organoids were then fixed in 4% paraformaldehyde for 15 min, permeabilized with 0.2% Triton X-100 for 1 h, and stained with DAPI (nuclei) and Alexa Fluor 647 phalloidin (actin) for 30 min. Images were acquired on an Olympus FV3000 laser scanning confocal microscope with a 30×/1.05 silicone objective. The voxel size was 0.414 × 0.414 × 3 µm, and both channels were imaged at the same time.

#### Cardiac tissue (confocal microscopy)

Formalin-fixed, paraffin-embedded heart sections (16 μm) were mounted on glass slides and deparaffinized. To reduce nonspecific binding, sections were blocked for 30 min at room temperature in 3% bovine serum albumin (BSA; 3737.3, Carl Roth) in Hanks’ Balanced Salt Solution (HBSS; 14025-100, Thermo Fisher). Sections were then stained with wheat germ agglutinin (WGA; 1:80, BOT-29026-1, Biozol), DAPI, a rabbit anti-RPS9 primary antibody (1:100, PA5-104493, Invitrogen) and an Alexa Fluor 488-conjugated anti-rabbit secondary antibody (1:100, A11008, Invitrogen). Autofluorescence was quenched with the Vector TrueVIEW kit (VEC-SP-8500, Biozol) before mounting. Sections were imaged on an Olympus IXplore SpinSR super-resolution system (Evident) with a 60× objective.

#### Breast tissue (histopathology)

We used whole slide images (WSIs) of breast cancer tissue from the HISTAI dataset^173^, which pairs WSIs with the corresponding pathology reports. We selected Ki67-stained slides from cases that pathologists had classified as grade 3 (poorly differentiated, with little gland formation), since lower-grade tumors would have required an additional tissue segmentation to delineate tumor regions. We excluded cases for which the report contained no or multiple Ki67 indices, as well as slides with imaging or severe tissue artefacts. This resulted in 237 WSIs, each with a slide-level Ki67 index from the pathology report, which we used as reference.

#### Platynereis larva (live imaging, light sheet microscopy)

Zygotes of *Platynereis dumerilii* were micro-injected with 181 ng/µl of mRNA encoding for H-RAS-mScarlet (membrane tag)^221^, together with 294 ng/µl of mRNA encoding H2B-GFP, as nuclear marker. The embryo was embedded in 1.5 % low-melting-point agarose in filtered artificial sea water (FASW), in a fluorinated ethylene propylene (FEP) tube (0.8 mm I.D./1.2 mm O.D). The time-lapse movie of the developing embryo was recorded on a Flamingo light sheet fluorescence microscope^222^ with the sample chamber filled with FASW. The timelapse recording started at three hours post fertilisation with a time interval of two minutes. The microscope was equipped with two water immersion objective lenses for illumination (CFI P-Fluor 10×W/0.30/3.50; Nikon) and two water immersion objective lenses for detection (CFI-75 LWD 16×W/0.80/3,00; Nikon), acquiring two views separated by 180°. Image stacks captured with the two detection arms were then fused for each channel by average blending using an in-house python script. Finally, the drift correction of the entire time series stacks was performed using Fast4DReg^223^.

#### Organoids (electron microscopy)

Organoids were grown from hct116 cells, a colorectal cancer cell line, following a similar protocol as described for the breast organoids. Samples were fixed in 2.5% glutaraldehyde at 37 °C and processed using an enhanced heavy-metal staining protocol. Briefly, samples were sequentially treated with reduced osmium tetroxide, thiocarbohydrazide, osmium tetroxide, uranyl acetate and lead aspartate, followed by dehydration through a graded ethanol series and embedding in Araldite resin. Resin-embedded samples were trimmed to expose the region of interest and mounted onto aluminium specimen pins using conductive epoxy. The block surfaces were coated with a conductive material when required to minimise charging during imaging. Serial block-face imaging was performed using a JSM-IT710HR/LV SEM (JEOL) equipped with a Katana automated ultramicrotome (ConnectomX). Images were acquired using a backscattered electron detector at an accelerating voltage of 4 kV, a pixel size of 16.6 nm and a section thickness of 100 nm. A total of 950 serial sections were collected per volume. We used one representative sample out of multiple acquired volumes for the use-case.

## Data Availability

We used publicly available datasets for model training and evaluation. See **Ext. Data Table 2** for details. We generated new data for some of the user studies. This data will be made available upon request, pending publication of associated biological studies in some cases.

## Code Availability

The μSAM software is available on GitHub under a permissive open-source license: https://github.com/computational-cell-analytics/micro-sam. The current version at the time of submission of this manuscript is 2.0.0. This repository also contains all the scripts used for training and evaluating the models. The software’s documentation is available at https://computational-cell-analytics.github.io/micro-sam/micro_sam.html. The μSAM2 models are currently hosted on ownCloud and will soon be uploaded to BioImage.IO.

## Acknowledgements

We would like to thank Marei Freitag, Luca Freckmann, Maryam Raahemi, Antonio Politi and Vanessa Goldschmidt for beta-testing the μSAM2 tool. Donglai Wei for contributing a MICRONS crop with mitochondrion annotations, and group members of AB, DG and JH for contributions on data acquisition. Julian Rüger for high performance cluster support. Further, we would like to thank Maryam Raahemi, Cyril Achard, Benjamin Gallusser and Martin Weigert for discussions about automatic tracking; Fatma Müge Tekin for discussions about volumetric segmentation; Jan Friedrich Meier for interactive segmentation discussions; Albert Dominguez Mantes for discussions about promptable segmentation; and Alexander Ecker for general discussions about methodology and offering Macbook for software testing and development. We would like to thank the developers of napari, Zarr, OME-NGFF, Trackastra, Traccuracy, Cell Tracking Challenge (CTC), GEFF, and segment-anything-2 for open-sourcing their codebase.

The work of AA was funded by the Deutsche Forschungsgemeinschaft (DFG, German Research Foundation) - PA 4341/2-1. This work is supported by the Ministry of Science and Culture of Lower Saxony through funds from the program zukunft.niedersachsen of the Volkswagen Foundation for the ‘CAIMed – Lower Saxony Center for Artificial Intelligence and Causal Methods in Medicine’ project (grant no. ZN4257). It was also supported by the Google Research Scholarship “Vision Foundation Models for Bioimage Segmentation”. The authors gratefully acknowledge the computing time granted by the KISSKI project and the Resource Allocation Board and provided on the supercomputer Emmy/Grete at NHR-Nord@Göttingen as part of the NHR infrastructure. The calculations for this research were conducted with computing resources under the projects kisski_usam2_pape and nim00007. The work of LM, NV and DG was funded by the Deutsche Forschungsgemeinschaft (DFG, German Research Foundation) - SFB1525 (INST 93/1204-1) and TRR 221/3 (INST 89/681-1).

## Author Contributions

CP and AA conceived the project, CP supervised it. AA and CP designed and implemented the μSAM2 model architecture and training procedure. AA curated and processed the training and evaluation datasets. AA, CT, TG, and CP implemented the interactive segmentation, classification and tracking functionality. AA developed and trained the dedicated models for electron microscopy. AA and CT performed the segmentation, classification and tracking evaluations. AA, TG, LM, BL, MA, NV, MH-T, IB, DG, AB, JH, and CP carried out the use-case applications and contributed biological data and domain expertise. AA and SvH visualized the results and designed the figures. AA and CP wrote the manuscript with input from all authors. All authors read and approved the final manuscript.

## Extended data figures and tables

**Extended Data Figure 1:**
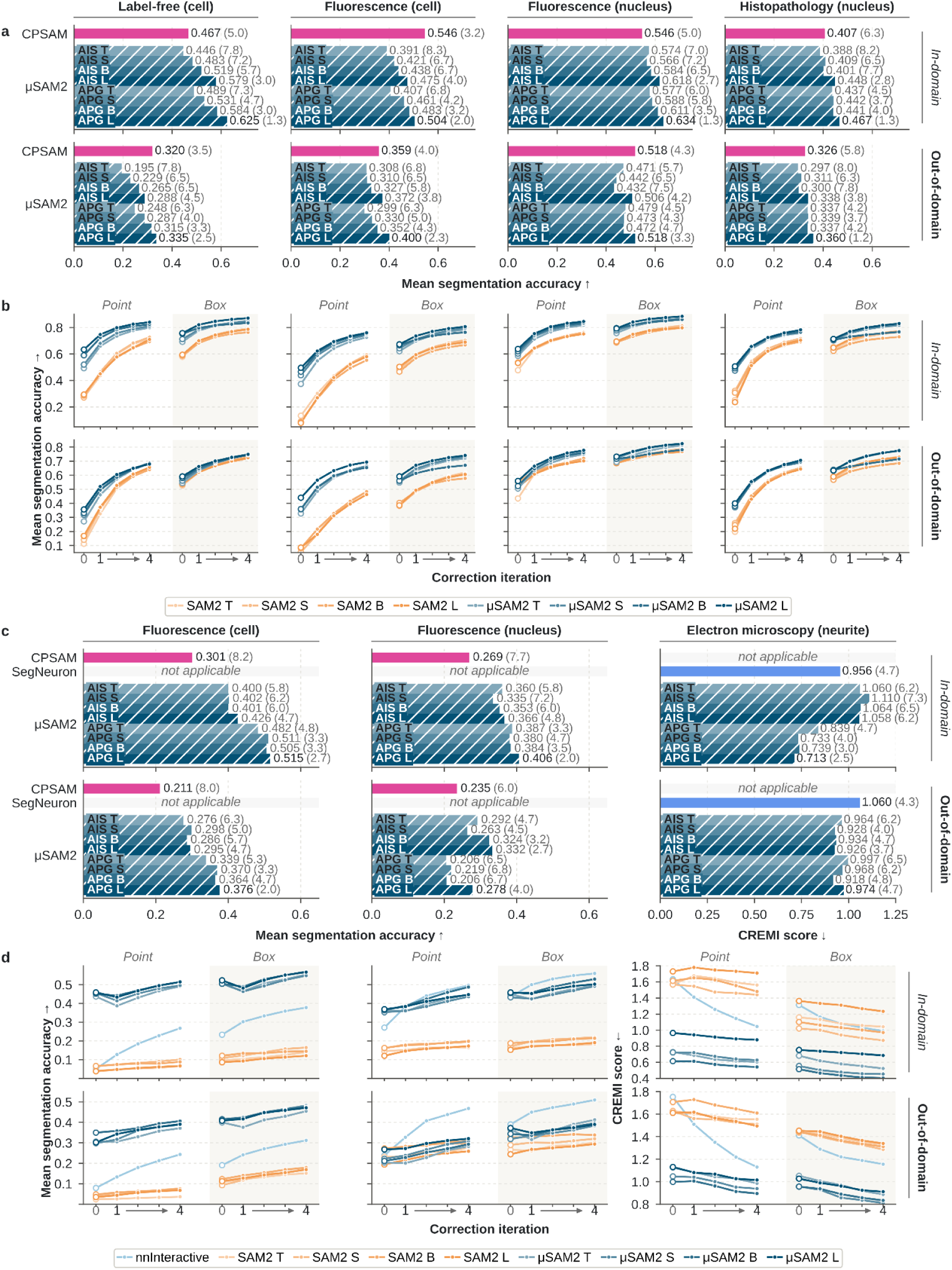
Segmentation with different model sizes. **a.** Automatic 2D segmentation with μSAM2 (AIS and APG) for different model sizes (T: hvit_t_, S: hvit_s_, B: hvit_b_, L: hvit_l_). Same set-up as Fig. 2 a. **b.** Interactive 2D segmentation with different model sizes. Same set-up as Fig. 2 c. **c.** Automatic 3D segmentation for the different model sizes. Same set-up as Fig. 3 a. **d.** Interactive 3D segmentation for different model sizes. Same setup as Fig. 3c.

**Extended Data Figure 2:**
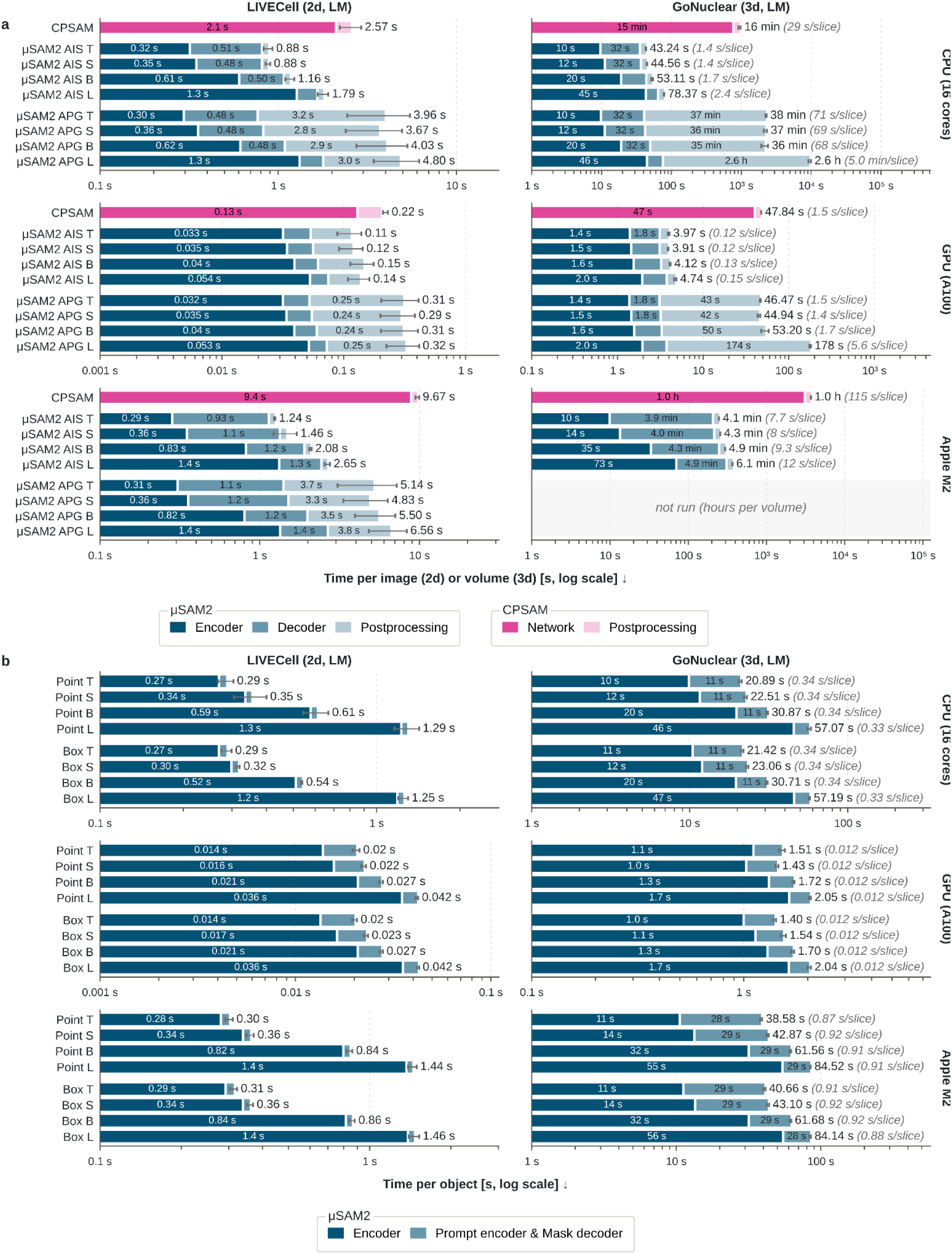
Segmentation runtimes. **a.** Runtimes for automatic segmentation (2D and 3D). We distinguish the runtime of the encoder, the segmentation decoder, and post-processing for μSAM2, and the network and post-processing for CellposeSAM. **b.** Runtimes for interactive segmentation (2D and 3D). We report the time for the image encoder and combined time for the prompt encoder and mask decoder. The image encoder has to be applied only once per image. We use a logarithmic scale to show disparate time measurements; hence, bar lengths are not comparable. GPU and CPU use bf16 precision, whereas the M processor uses fp32 due to a missing bf16 kernel. This explains why it is not faster than the CPU. We will implement this kernel soon.

**Extended Data Figure 3:**
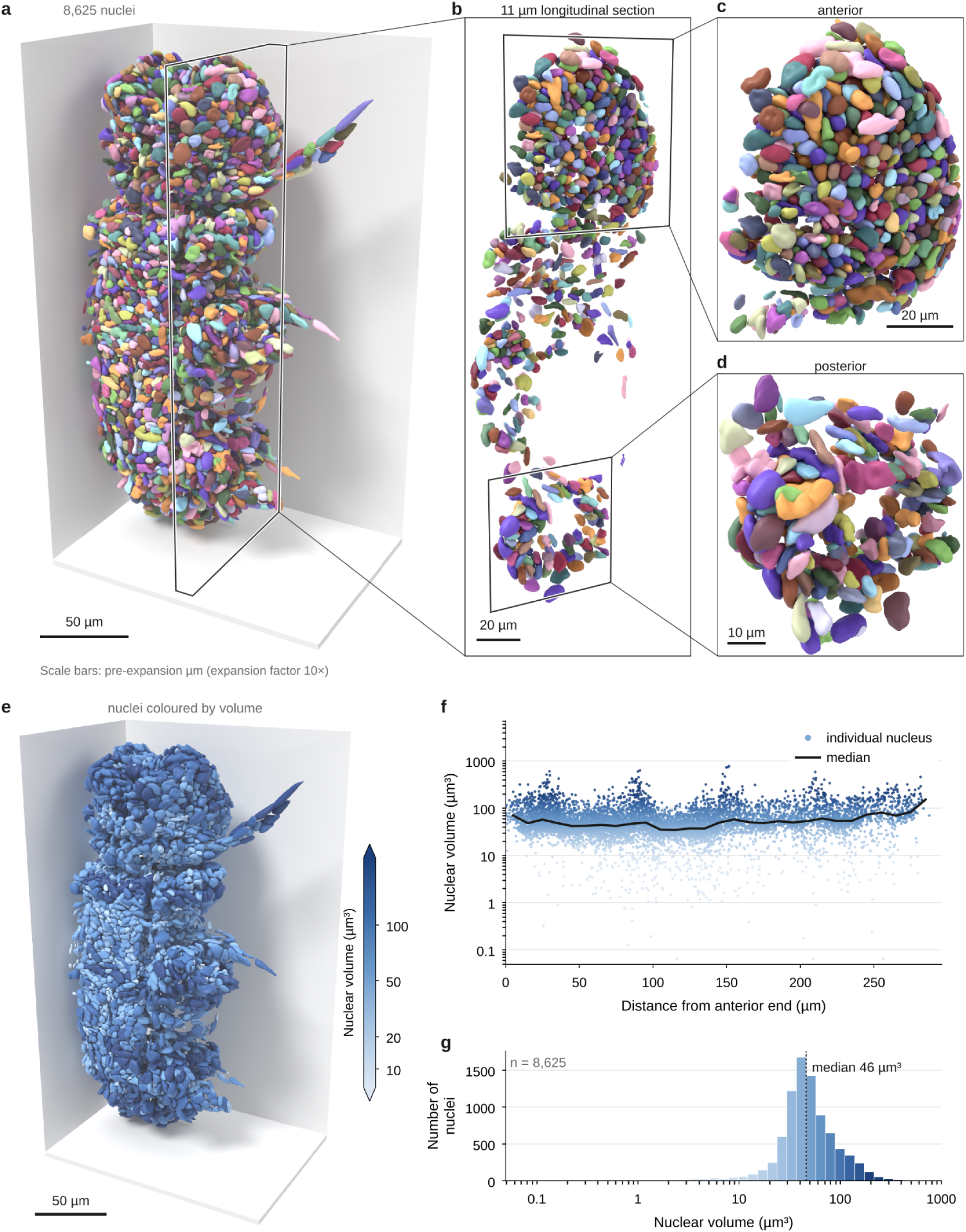
Segmentation and analysis of a *P. dumerilii larva*. a–c. Nuclei of a *P. dumerilii* larva imaged in a light-sheet microscope after expansion, segmented with μSAM2 (AIS), rendered in different views and zoom-ins. **e.** Nucleus segmentation color-coded by the measured nuclear volume. **f.** Nuclear volume across the anterior axis. **g.** Histogram of the nuclear volume.

**Extended Data Figure 4:**
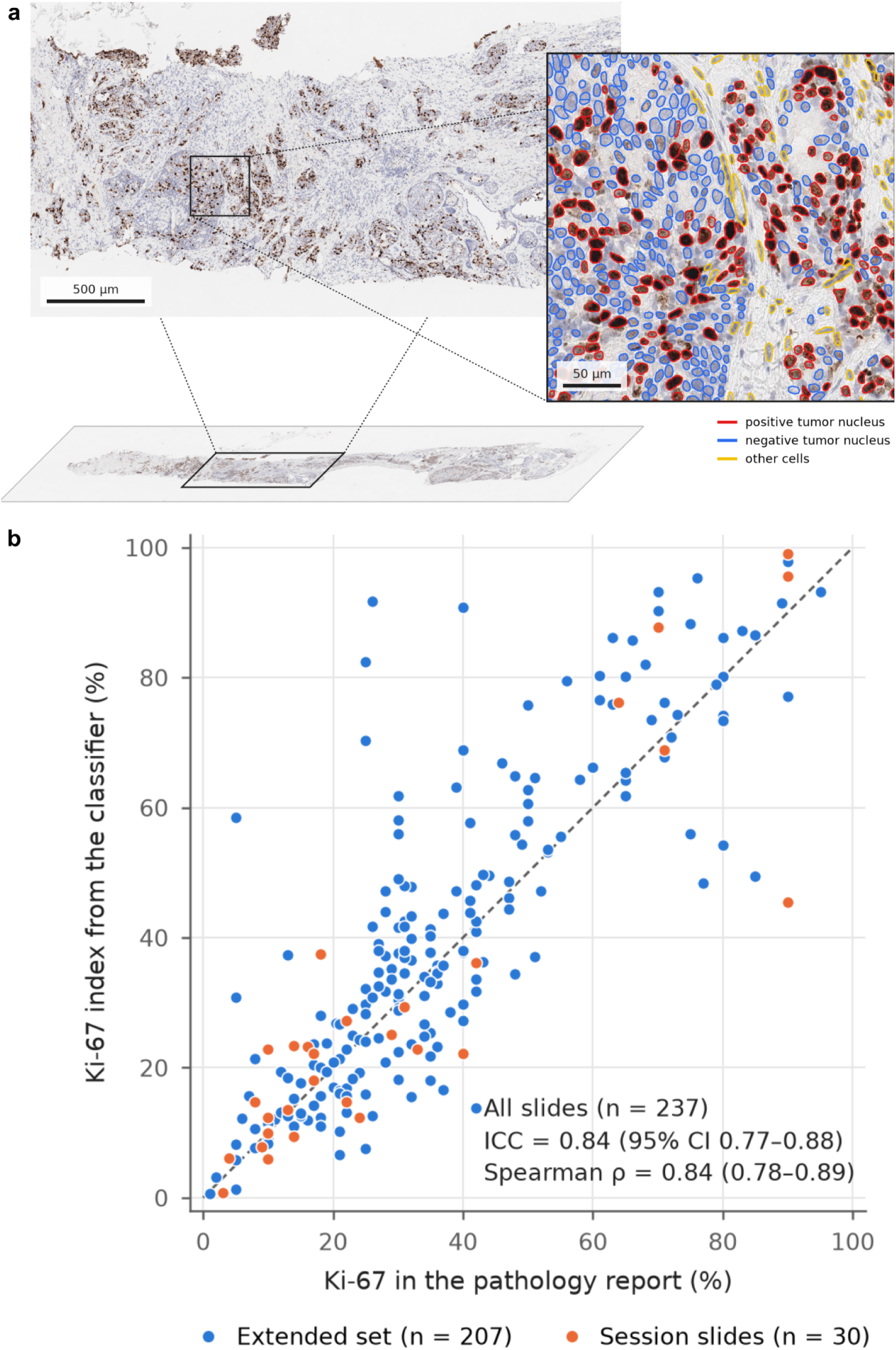
Nucleus segmentation and classification for tumor scoring. **a.** Example nucleus segmentation and classification for a WSI with DAB stain, from μSAM2 automatic segmentation (APG) and a classifier trained via the object classification widget and then applied automatically to all segmented nuclei. **b.** Correlation of the Ki67 index derived with our approach and the pathologist scoring from HISTAI^173^. “Session slides” indicate the WSIs used to train the classifier.

**Extended Data Table 1:**
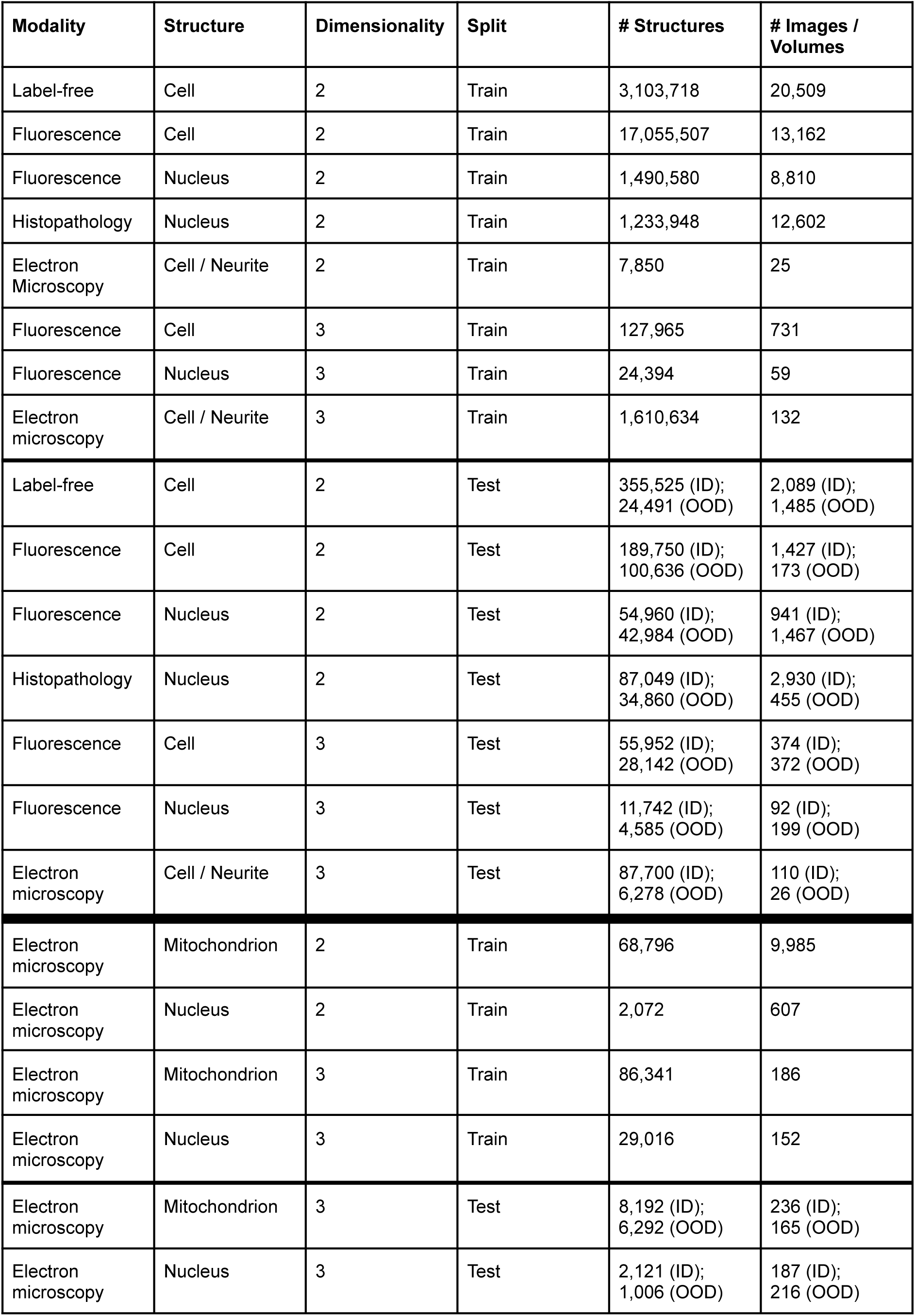
Overview of the dataset sizes differentiated by imaging modality, labeled structure, and data dimensionality, and data split (training or test data).

**Extended Data Table 2:**
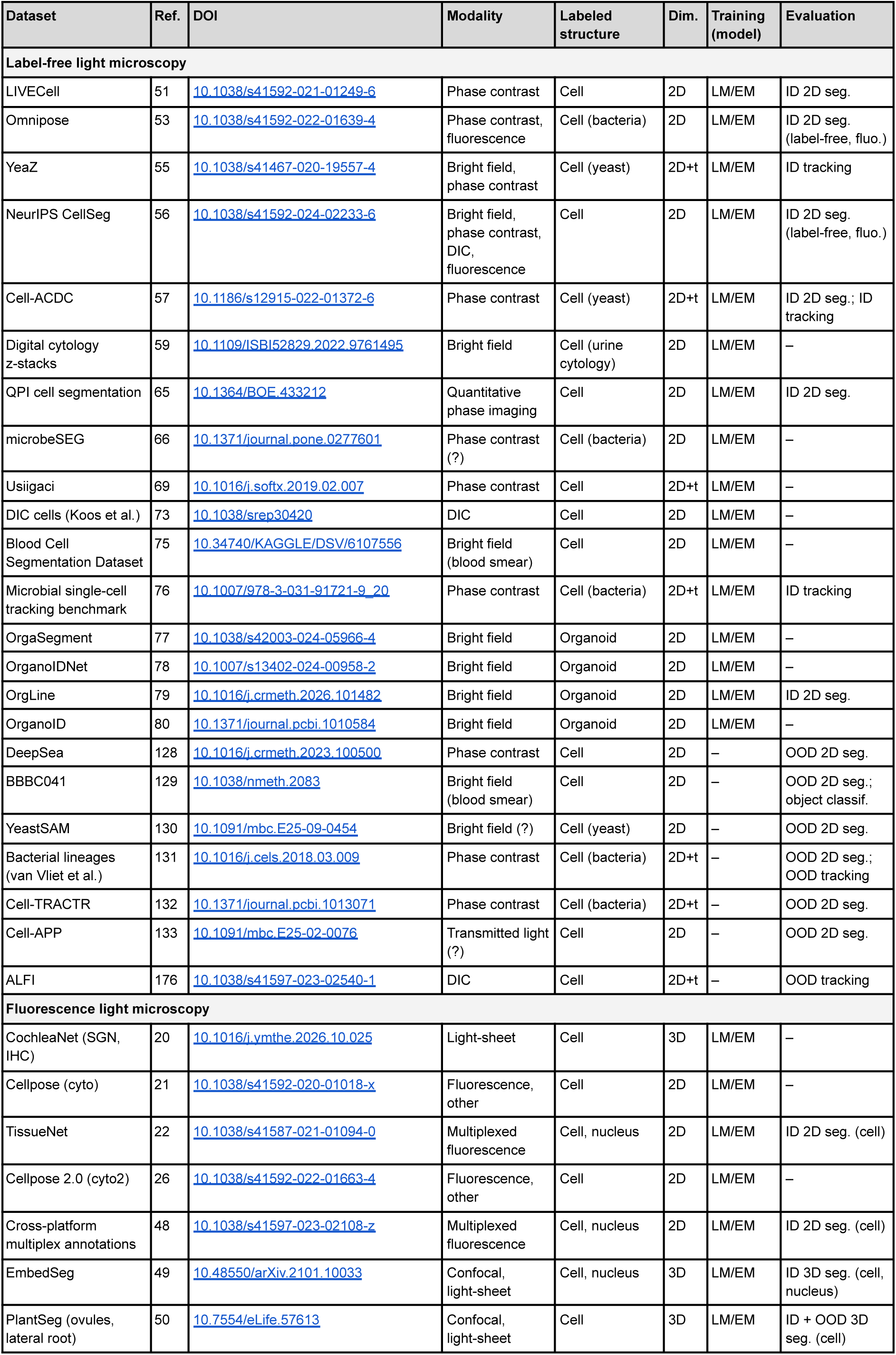
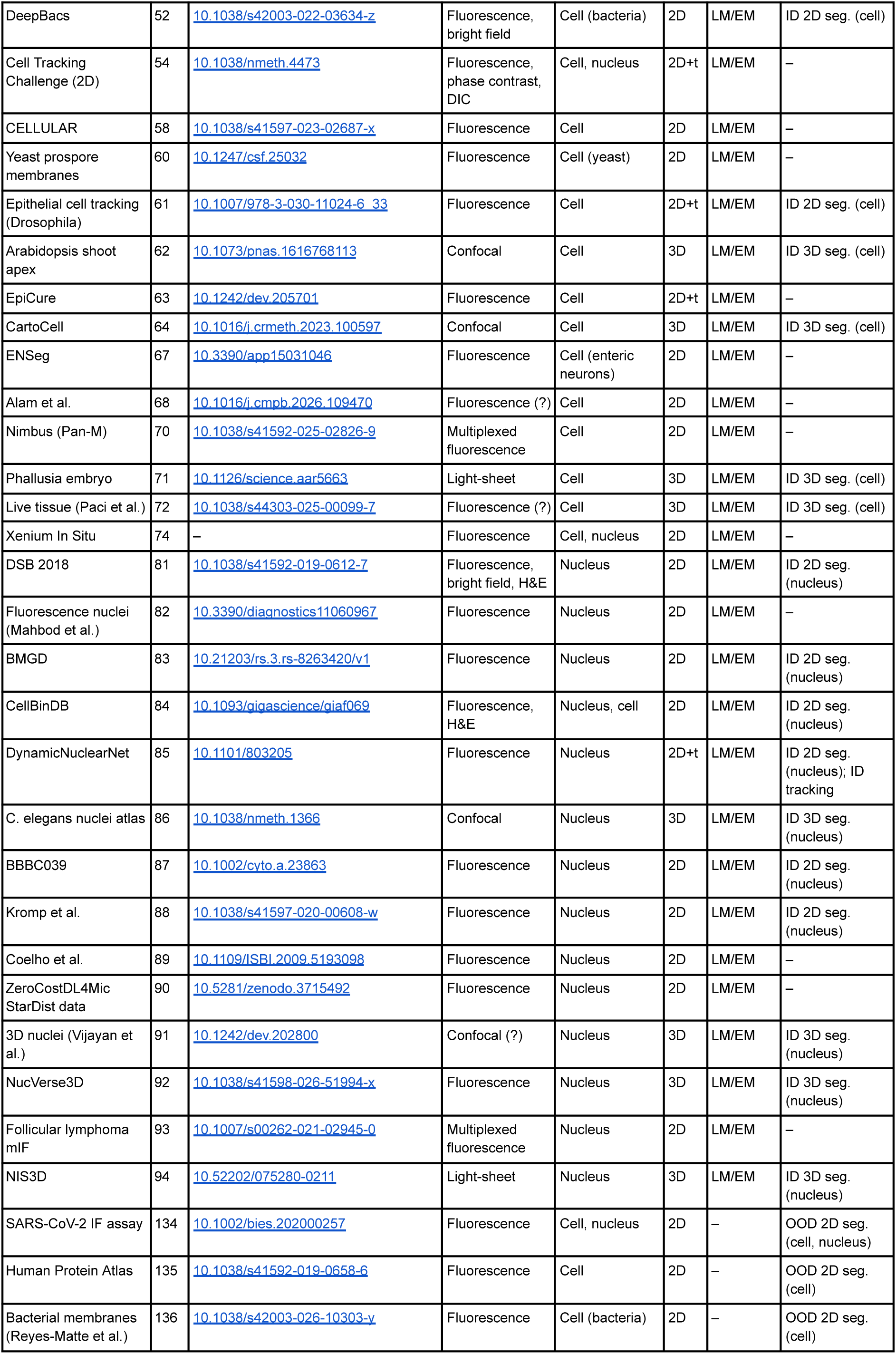
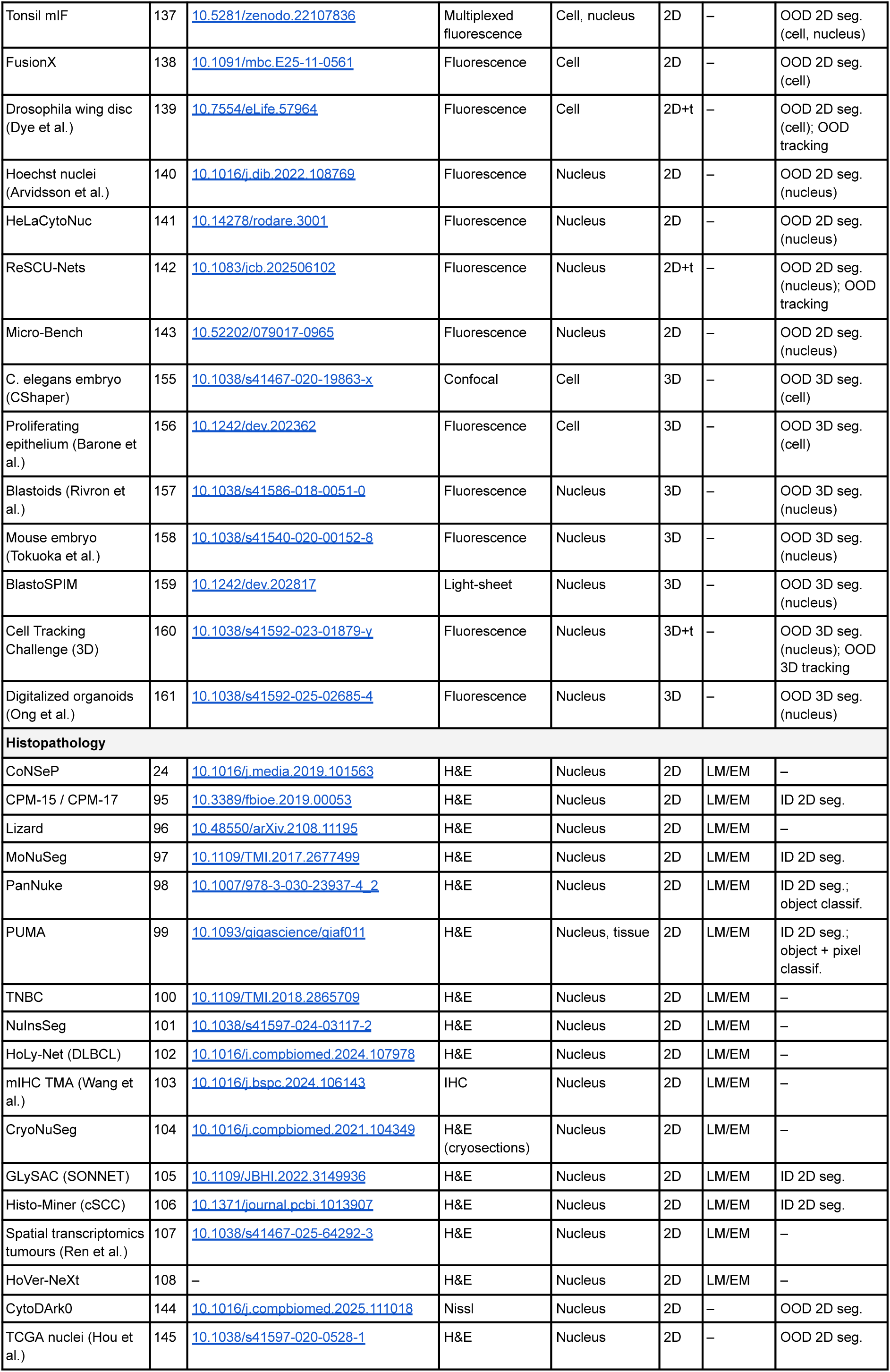
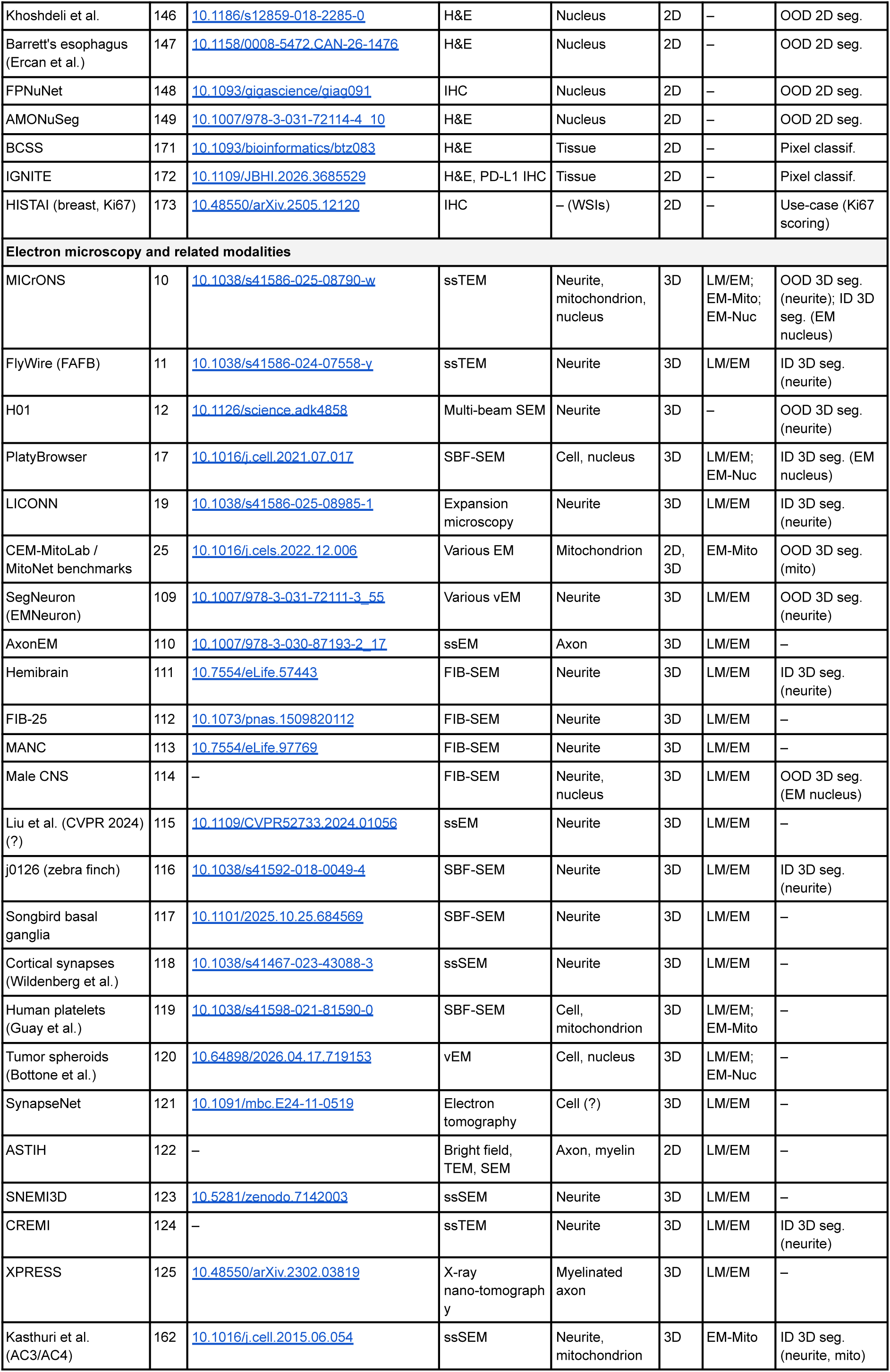
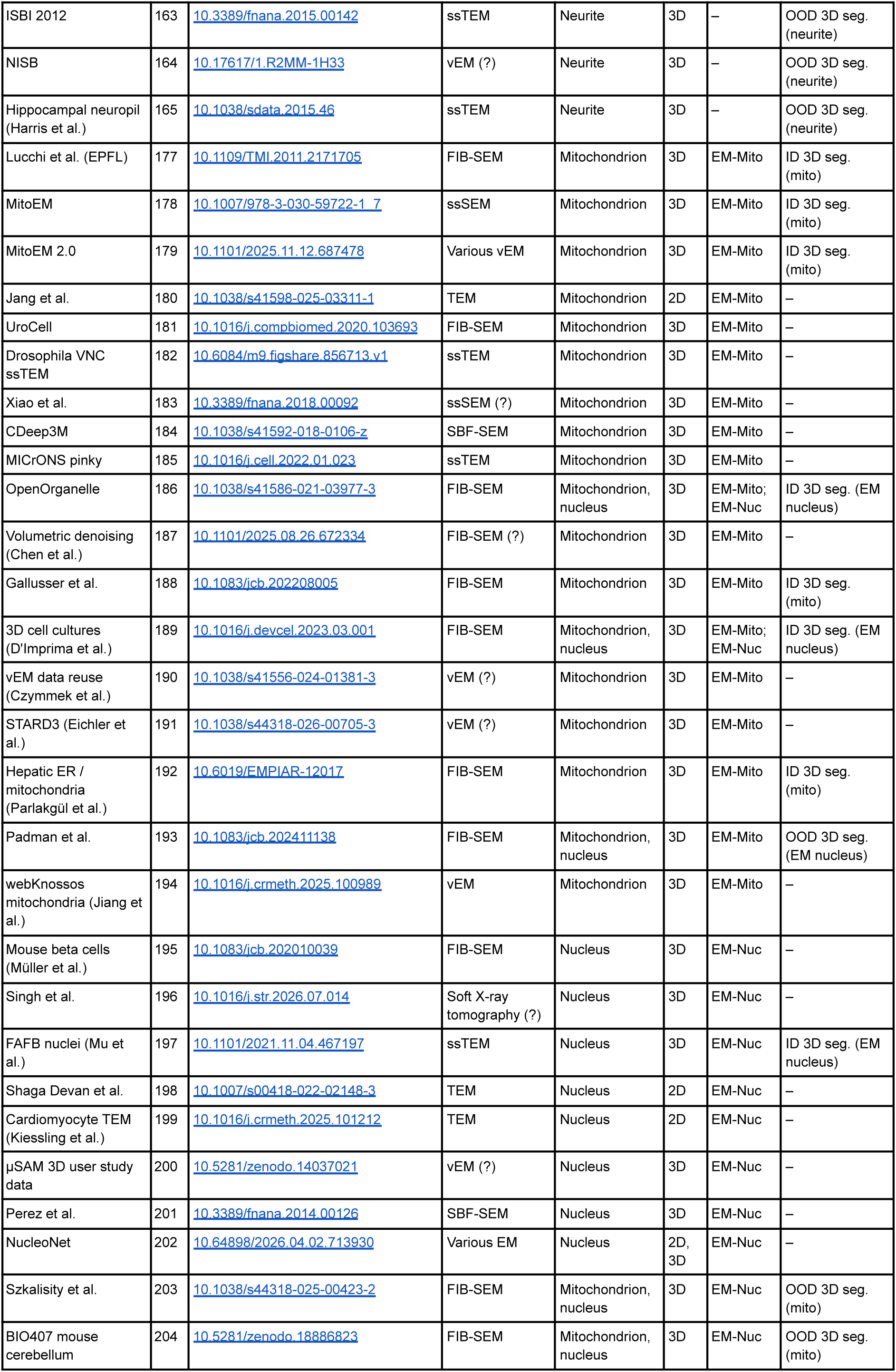
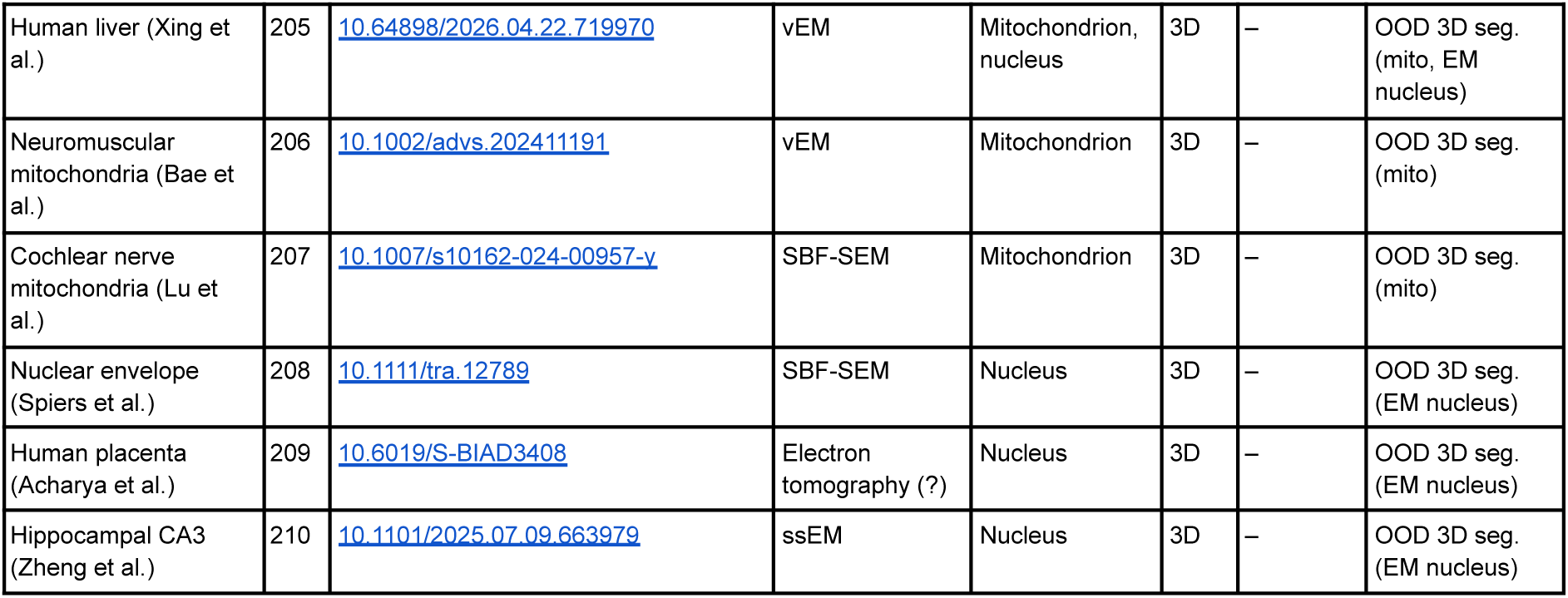
List of all datasets used in this study.

| Dataset | Ref. | DOI | Modality | Labeled structure | Dim. | Training (model) | Evaluation |
|---|---|---|---|---|---|---|---|
| **Label-free light microscopy** |  |  |  |  |  |  |  |
| LIVECell | 51 | 10.1038/s41592-021-01249-6 | Phase contrast | Cell | 2D | LM/EM | ID 2D seg. |
| Omnipose | 53 | 10.1038/s41592-022-01639-4 | Phase contrast, fluorescence | Cell (bacteria) | 2D | LM/EM | ID 2D seg. (label-free, fluo.) |
| YeaZ | 55 | 10.1038/s41467-020-19557-4 | Bright field, phase contrast | Cell (yeast) | 2D+t | LM/EM | ID tracking |
| NeurIPS CellSeg | 56 | 10.1038/s41592-024-02233-6 | Bright field, phase contrast, DIC, fluorescence | Cell | 2D | LM/EM | ID 2D seg. (label-free, fluo.) |
| Cell-ACDC | 57 | 10.1186/s12915-022-01372-6 | Phase contrast | Cell (yeast) | 2D+t | LM/EM | ID 2D seg.; ID tracking |
| Digital cytology z-stacks | 59 | 10.1109/ISBI52829.2022.9761495 | Bright field | Cell (urine cytology) | 2D | LM/EM | – |
| QPI cell segmentation | 65 | 10.1364/BOE.433212 | Quantitative phase imaging | Cell | 2D | LM/EM | ID 2D seg. |
| microbeSEG | 66 | 10.1371/journal.pone.0277601 | Phase contrast (?) | Cell (bacteria) | 2D | LM/EM | – |
| Usiigaci | 69 | 10.1016/j.softx.2019.02.007 | Phase contrast | Cell | 2D+t | LM/EM | – |
| DIC cells (Koos et al.) | 73 | 10.1038/srep30420 | DIC | Cell | 2D | LM/EM | – |
| Blood Cell Segmentation Dataset | 75 | 10.34740/KAGGLE/DSV/6107556 | Bright field (blood smear) | Cell | 2D | LM/EM | – |
| Microbial single-cell tracking benchmark | 76 | 10.1007/978-3-031-91721-9_20 | Phase contrast | Cell (bacteria) | 2D+t | LM/EM | ID tracking |
| OrgaSegment | 77 | 10.1038/s42003-024-05966-4 | Bright field | Organoid | 2D | LM/EM | – |
| OrganoIDNet | 78 | 10.1007/s13402-024-00958-2 | Bright field | Organoid | 2D | LM/EM | – |
| OrgLine | 79 | 10.1016/j.crmeth.2026.101482 | Bright field | Organoid | 2D | LM/EM | ID 2D seg. |
| OrganoID | 80 | 10.1371/journal.pcbi.1010584 | Bright field | Organoid | 2D | LM/EM | – |
| DeepSea | 128 | 10.1016/j.crmeth.2023.100500 | Phase contrast | Cell | 2D | – | OOD 2D seg. |
| BBBC041 | 129 | 10.1038/nmeth.2083 | Bright field (blood smear) | Cell | 2D | – | OOD 2D seg.; object classif. |
| YeastSAM | 130 | 10.1091/mbc.E25-09-0454 | Bright field (?) | Cell (yeast) | 2D | – | OOD 2D seg. |
| Bacterial lineages (van Vliet et al.) | 131 | 10.1016/j.cels.2018.03.009 | Phase contrast | Cell (bacteria) | 2D+t | – | OOD 2D seg.; OOD tracking |
| Cell-TRACTR | 132 | 10.1371/journal.pcbi.1013071 | Phase contrast | Cell (bacteria) | 2D+t | – | OOD 2D seg. |
| Cell-APP | 133 | 10.1091/mbc.E25-02-0076 | Transmitted light (?) | Cell | 2D | – | OOD 2D seg. |
| ALFI | 176 | 10.1038/s41597-023-02540-1 | DIC | Cell | 2D+t | – | OOD tracking |
| **Fluorescence light microscopy** |  |  |  |  |  |  |  |
| CochleaNet (SGN, IHC) | 20 | 10.1016/j.ymthe.2026.10.025 | Light-sheet | Cell | 3D | LM/EM | – |
| Cellpose (cyto) | 21 | 10.1038/s41592-020-01018-x | Fluorescence, other | Cell | 2D | LM/EM | – |
| TissueNet | 22 | 10.1038/s41587-021-01094-0 | Multiplexed fluorescence | Cell, nucleus | 2D | LM/EM | ID 2D seg. (cell) |
| Cellpose 2.0 (cyto2) | 26 | 10.1038/s41592-022-01663-4 | Fluorescence, other | Cell | 2D | LM/EM | – |
| Cross-platform multiplex annotations | 48 | 10.1038/s41597-023-02108-z | Multiplexed fluorescence | Cell, nucleus | 2D | LM/EM | ID 2D seg. (cell) |
| EmbedSeg | 49 | 10.48550/arXiv.2101.10033 | Confocal, light-sheet | Cell, nucleus | 3D | LM/EM | ID 3D seg. (cell, nucleus) |
| PlantSeg (ovules, lateral root) | 50 | 10.7554/eLife.57613 | Confocal, light-sheet | Cell | 3D | LM/EM | ID + OOD 3D seg. (cell) |
| DeepBacs | 52 | 10.1038/s42003-022-03634-z | Fluorescence, bright field | Cell (bacteria) | 2D | LM/EM | ID 2D seg. (cell) |
| Cell Tracking Challenge (2D) | 54 | 10.1038/nmeth.4473 | Fluorescence, phase contrast, DIC | Cell, nucleus | 2D+t | LM/EM | – |
| CELLULAR | 58 | 10.1038/s41597-023-02687-x | Fluorescence | Cell | 2D | LM/EM | – |
| Yeast prospore membranes | 60 | 10.1247/csf.25032 | Fluorescence | Cell (yeast) | 2D | LM/EM | – |
| Epithelial cell tracking (Drosophila) | 61 | 10.1007/978-3-030-11024-6_33 | Fluorescence | Cell | 2D+t | LM/EM | ID 2D seg. (cell) |
| Arabidopsis shoot apex | 62 | 10.1073/pnas.1616768113 | Confocal | Cell | 3D | LM/EM | ID 3D seg. (cell) |
| EpiCure | 63 | 10.1242/dev.205701 | Fluorescence | Cell | 2D+t | LM/EM | – |
| CartoCell | 64 | 10.1016/j.crmeth.2023.100597 | Confocal | Cell | 3D | LM/EM | ID 3D seg. (cell) |
| ENSeg | 67 | 10.3390/app15031046 | Fluorescence | Cell (enteric neurons) | 2D | LM/EM | – |
| Alam et al. | 68 | 10.1016/j.cmpb.2026.109470 | Fluorescence (?) | Cell | 2D | LM/EM | – |
| Nimbus (Pan-M) | 70 | 10.1038/s41592-025-02826-9 | Multiplexed fluorescence | Cell | 2D | LM/EM | – |
| Phallusia embryo | 71 | 10.1126/science.aar5663 | Light-sheet | Cell | 3D | LM/EM | ID 3D seg. (cell) |
| Live tissue (Paci et al.) | 72 | 10.1038/s44303-025-00099-7 | Fluorescence (?) | Cell | 3D | LM/EM | ID 3D seg. (cell) |
| Xenium In Situ | 74 | – | Fluorescence | Cell, nucleus | 2D | LM/EM | – |
| DSB 2018 | 81 | 10.1038/s41592-019-0612-7 | Fluorescence, bright field, H&E | Nucleus | 2D | LM/EM | ID 2D seg. (nucleus) |
| Fluorescence nuclei (Mahbod et al.) | 82 | 10.3390/diagnostics11060967 | Fluorescence | Nucleus | 2D | LM/EM | – |
| BMGD | 83 | 10.21203/rs.3.rs-8263420/v1 | Fluorescence | Nucleus | 2D | LM/EM | ID 2D seg. (nucleus) |
| CellBinDB | 84 | 10.1093/gigascience/giaf069 | Fluorescence, H&E | Nucleus, cell | 2D | LM/EM | ID 2D seg. (nucleus) |
| DynamicNuclearNet | 85 | 10.1101/803205 | Fluorescence | Nucleus | 2D+t | LM/EM | ID 2D seg. (nucleus); ID tracking |
| C. elegans nuclei atlas | 86 | 10.1038/nmeth.1366 | Confocal | Nucleus | 3D | LM/EM | ID 3D seg. (nucleus) |
| BBBC039 | 87 | 10.1002/cyto.a.23863 | Fluorescence | Nucleus | 2D | LM/EM | ID 2D seg. (nucleus) |
| Kromp et al. | 88 | 10.1038/s41597-020-00608-w | Fluorescence | Nucleus | 2D | LM/EM | ID 2D seg. (nucleus) |
| Coelho et al. | 89 | 10.1109/ISBI.2009.5193098 | Fluorescence | Nucleus | 2D | LM/EM | – |
| ZeroCostDL4Mic StarDist data | 90 | 10.5281/zenodo.3715492 | Fluorescence | Nucleus | 2D | LM/EM | – |
| 3D nuclei (Vijayan et al.) | 91 | 10.1242/dev.202800 | Confocal (?) | Nucleus | 3D | LM/EM | ID 3D seg. (nucleus) |
| NucVerse3D | 92 | 10.1038/s41598-026-51994-x | Fluorescence | Nucleus | 3D | LM/EM | ID 3D seg. (nucleus) |
| Follicular lymphoma mIF | 93 | 10.1007/s00262-021-02945-0 | Multiplexed fluorescence | Nucleus | 2D | LM/EM | – |
| NIS3D | 94 | 10.52202/075280-0211 | Light-sheet | Nucleus | 3D | LM/EM | ID 3D seg. (nucleus) |
| SARS-CoV-2 IF assay | 134 | 10.1002/bies.202000257 | Fluorescence | Cell, nucleus | 2D | – | OOD 2D seg. (cell, nucleus) |
| Human Protein Atlas | 135 | 10.1038/s41592-019-0658-6 | Fluorescence | Cell | 2D | – | OOD 2D seg. (cell) |
| Bacterial membranes (Reyes-Matte et al.) | 136 | 10.1038/s42003-026-10303-y | Fluorescence | Cell (bacteria) | 2D | – | OOD 2D seg. (cell) |
| Tonsil mIF | 137 | 10.5281/zenodo.22107836 | Multiplexed fluorescence | Cell, nucleus | 2D | – | OOD 2D seg. (cell, nucleus) |
| FusionX | 138 | 10.1091/mbc.E25-11-0561 | Fluorescence | Cell | 2D | – | OOD 2D seg. (cell) |
| Drosophila wing disc (Dye et al.) | 139 | 10.7554/eLife.57964 | Fluorescence | Cell | 2D+t | – | OOD 2D seg. (cell); OOD tracking |
| Hoechst nuclei (Arvidsson et al.) | 140 | 10.1016/j.dib.2022.108769 | Fluorescence | Nucleus | 2D | – | OOD 2D seg. (nucleus) |
| HeLaCytoNuc | 141 | 10.14278/rodare.3001 | Fluorescence | Nucleus | 2D | – | OOD 2D seg. (nucleus) |
| ReSCU-Nets | 142 | 10.1083/jcb.202506102 | Fluorescence | Nucleus | 2D+t | – | OOD 2D seg. (nucleus); OOD tracking |
| Micro-Bench | 143 | 10.52202/079017-0965 | Fluorescence | Nucleus | 2D | – | OOD 2D seg. (nucleus) |
| C. elegans embryo (CShaper) | 155 | 10.1038/s41467-020-19863-x | Confocal | Cell | 3D | – | OOD 3D seg. (cell) |
| Proliferating epithelium (Barone et al.) | 156 | 10.1242/dev.202362 | Fluorescence | Cell | 3D | – | OOD 3D seg. (cell) |
| Blastoids (Rivron et al.) | 157 | 10.1038/s41586-018-0051-0 | Fluorescence | Nucleus | 3D | – | OOD 3D seg. (nucleus) |
| Mouse embryo (Tokuoka et al.) | 158 | 10.1038/s41540-020-00152-8 | Fluorescence | Nucleus | 3D | – | OOD 3D seg. (nucleus) |
| BlastoSPIM | 159 | 10.1242/dev.202817 | Light-sheet | Nucleus | 3D | – | OOD 3D seg. (nucleus) |
| Cell Tracking Challenge (3D) | 160 | 10.1038/s41592-023-01879-y | Fluorescence | Nucleus | 3D+t | – | OOD 3D seg. (nucleus); OOD 3D tracking |
| Digitalized organoids (Ong et al.) | 161 | 10.1038/s41592-025-02685-4 | Fluorescence | Nucleus | 3D | – | OOD 3D seg. (nucleus) |
| **Histopathology** |  |  |  |  |  |  |  |
| CoNSeP | 24 | 10.1016/j.media.2019.101563 | H&E | Nucleus | 2D | LM/EM | – |
| CPM-15 / CPM-17 | 95 | 10.3389/fbioe.2019.00053 | H&E | Nucleus | 2D | LM/EM | ID 2D seg. |
| Lizard | 96 | 10.48550/arXiv.2108.11195 | H&E | Nucleus | 2D | LM/EM | – |
| MoNuSeg | 97 | 10.1109/TMI.2017.2677499 | H&E | Nucleus | 2D | LM/EM | ID 2D seg. |
| PanNuke | 98 | 10.1007/978-3-030-23937-4_2 | H&E | Nucleus | 2D | LM/EM | ID 2D seg.; object classif. |
| PUMA | 99 | 10.1093/gigascience/giaf011 | H&E | Nucleus, tissue | 2D | LM/EM | ID 2D seg.; object + pixel classif. |
| TNBC | 100 | 10.1109/TMI.2018.2865709 | H&E | Nucleus | 2D | LM/EM | – |
| NuInsSeg | 101 | 10.1038/s41597-024-03117-2 | H&E | Nucleus | 2D | LM/EM | – |
| HoLy-Net (DLBCL) | 102 | 10.1016/j.compbiomed.2024.107978 | H&E | Nucleus | 2D | LM/EM | – |
| mIHC TMA (Wang et al.) | 103 | 10.1016/j.bspc.2024.106143 | IHC | Nucleus | 2D | LM/EM | – |
| CryoNuSeg | 104 | 10.1016/j.compbiomed.2021.104349 | H&E (cryosections) | Nucleus | 2D | LM/EM | – |
| GLySAC (SONNET) | 105 | 10.1109/JBHI.2022.3149936 | H&E | Nucleus | 2D | LM/EM | ID 2D seg. |
| Histo-Miner (cSCC) | 106 | 10.1371/journal.pcbi.1013907 | H&E | Nucleus | 2D | LM/EM | ID 2D seg. |
| Spatial transcriptomics tumours (Ren et al.) | 107 | 10.1038/s41467-025-64292-3 | H&E | Nucleus | 2D | LM/EM | – |
| HoVer-NeXt | 108 | – | H&E | Nucleus | 2D | LM/EM | – |
| CytoDArk0 | 144 | 10.1016/j.compbiomed.2025.111018 | Nissl | Nucleus | 2D | – | OOD 2D seg. |
| TCGA nuclei (Hou et al.) | 145 | 10.1038/s41597-020-0528-1 | H&E | Nucleus | 2D | – | OOD 2D seg. |
| Khoshdeli et al. | 146 | 10.1186/s12859-018-2285-0 | H&E | Nucleus | 2D | – | OOD 2D seg. |
| Barrett's esophagus (Ercan et al.) | 147 | 10.1158/0008-5472.CAN-26-1476 | H&E | Nucleus | 2D | – | OOD 2D seg. |
| FPNuNet | 148 | 10.1093/gigascience/giag091 | IHC | Nucleus | 2D | – | OOD 2D seg. |
| AMONuSeg | 149 | 10.1007/978-3-031-72114-4_10 | H&E | Nucleus | 2D | – | OOD 2D seg. |
| BCSS | 171 | 10.1093/bioinformatics/btz083 | H&E | Tissue | 2D | – | Pixel classif. |
| IGNITE | 172 | 10.1109/JBHI.2026.3685529 | H&E, PD-L1 IHC | Tissue | 2D | – | Pixel classif. |
| HISTAI (breast, Ki67) | 173 | 10.48550/arXiv.2505.12120 | IHC | – (WSIs) | 2D | – | Use-case (Ki67 scoring) |
| **Electron microscopy and related modalities** |  |  |  |  |  |  |  |
| MICrONS | 10 | 10.1038/s41586-025-08790-w | ssTEM | Neurite, mitochondrion, nucleus | 3D | LM/EM; EM-Mito; EM-Nuc | OOD 3D seg. (neurite); ID 3D seg. (EM nucleus) |
| FlyWire (FAFB) | 11 | 10.1038/s41586-024-07558-y | ssTEM | Neurite | 3D | LM/EM | ID 3D seg. (neurite) |
| H01 | 12 | 10.1126/science.adk4858 | Multi-beam SEM | Neurite | 3D | – | OOD 3D seg. (neurite) |
| PlatyBrowser | 17 | 10.1016/j.cell.2021.07.017 | SBF-SEM | Cell, nucleus | 3D | LM/EM; EM-Nuc | ID 3D seg. (EM nucleus) |
| LICONN | 19 | 10.1038/s41586-025-08985-1 | Expansion microscopy | Neurite | 3D | LM/EM | ID 3D seg. (neurite) |
| CEM-MitoLab / MitoNet benchmarks | 25 | 10.1016/j.cels.2022.12.006 | Various EM | Mitochondrion | 2D, 3D | EM-Mito | OOD 3D seg. (mito) |
| SegNeuron (EMNeuron) | 109 | 10.1007/978-3-031-72111-3_55 | Various vEM | Neurite | 3D | LM/EM | OOD 3D seg. (neurite) |
| AxonEM | 110 | 10.1007/978-3-030-87193-2_17 | ssEM | Axon | 3D | LM/EM | – |
| Hemibrain | 111 | 10.7554/eLife.57443 | FIB-SEM | Neurite | 3D | LM/EM | ID 3D seg. (neurite) |
| FIB-25 | 112 | 10.1073/pnas.1509820112 | FIB-SEM | Neurite | 3D | LM/EM | – |
| MANC | 113 | 10.7554/eLife.97769 | FIB-SEM | Neurite | 3D | LM/EM | – |
| Male CNS | 114 | – | FIB-SEM | Neurite, nucleus | 3D | LM/EM | OOD 3D seg. (EM nucleus) |
| Liu et al. (CVPR 2024) (?) | 115 | 10.1109/CVPR52733.2024.01056 | ssEM | Neurite | 3D | LM/EM | – |
| j0126 (zebra finch) | 116 | 10.1038/s41592-018-0049-4 | SBF-SEM | Neurite | 3D | LM/EM | ID 3D seg. (neurite) |
| Songbird basal ganglia | 117 | 10.1101/2025.10.25.684569 | SBF-SEM | Neurite | 3D | LM/EM | – |
| Cortical synapses (Wildenberg et al.) | 118 | 10.1038/s41467-023-43088-3 | ssSEM | Neurite | 3D | LM/EM | – |
| Human platelets (Guay et al.) | 119 | 10.1038/s41598-021-81590-0 | SBF-SEM | Cell, mitochondrion | 3D | LM/EM; EM-Mito | – |
| Tumor spheroids (Bottone et al.) | 120 | 10.64898/2026.04.17.719153 | vEM | Cell, nucleus | 3D | LM/EM; EM-Nuc | – |
| SynapseNet | 121 | 10.1091/mbc.E24-11-0519 | Electron tomography | Cell (?) | 3D | LM/EM | – |
| ASTIH | 122 | – | Bright field, TEM, SEM | Axon, myelin | 2D | LM/EM | – |
| SNEMI3D | 123 | 10.5281/zenodo.7142003 | ssSEM | Neurite | 3D | LM/EM | – |
| CREMI | 124 | – | ssTEM | Neurite | 3D | LM/EM | ID 3D seg. (neurite) |
| XPRESS | 125 | 10.48550/arXiv.2302.03819 | X-ray nano-tomography | Myelinated axon | 3D | LM/EM | – |
| Kasthuri et al. (AC3/AC4) | 162 | 10.1016/j.cell.2015.06.054 | ssSEM | Neurite, mitochondrion | 3D | EM-Mito | ID 3D seg. (neurite, mito) |
| ISBI 2012 | 163 | 10.3389/fnana.2015.00142 | ssTEM | Neurite | 3D | – | OOD 3D seg. (neurite) |
| NISB | 164 | 10.17617/1.R2MM-1H33 | vEM (?) | Neurite | 3D | – | OOD 3D seg. (neurite) |
| Hippocampal neuropil (Harris et al.) | 165 | 10.1038/sdata.2015.46 | ssTEM | Neurite | 3D | – | OOD 3D seg. (neurite) |
| Lucchi et al. (EPFL) | 177 | 10.1109/TMI.2011.2171705 | FIB-SEM | Mitochondrion | 3D | EM-Mito | ID 3D seg. (mito) |
| MitoEM | 178 | 10.1007/978-3-030-59722-1_7 | ssSEM | Mitochondrion | 3D | EM-Mito | ID 3D seg. (mito) |
| MitoEM 2.0 | 179 | 10.1101/2025.11.12.687478 | Various vEM | Mitochondrion | 3D | EM-Mito | ID 3D seg. (mito) |
| Jang et al. | 180 | 10.1038/s41598-025-03311-1 | TEM | Mitochondrion | 2D | EM-Mito | – |
| UroCell | 181 | 10.1016/j.compbiomed.2020.103693 | FIB-SEM | Mitochondrion | 3D | EM-Mito | – |
| Drosophila VNC ssTEM | 182 | 10.6084/m9.figshare.856713.v1 | ssTEM | Mitochondrion | 3D | EM-Mito | – |
| Xiao et al. | 183 | 10.3389/fnana.2018.00092 | ssSEM (?) | Mitochondrion | 3D | EM-Mito | – |
| CDeep3M | 184 | 10.1038/s41592-018-0106-z | SBF-SEM | Mitochondrion | 3D | EM-Mito | – |
| MICrONS pinky | 185 | 10.1016/j.cell.2022.01.023 | ssTEM | Mitochondrion | 3D | EM-Mito | – |
| OpenOrganelle | 186 | 10.1038/s41586-021-03977-3 | FIB-SEM | Mitochondrion, nucleus | 3D | EM-Mito; EM-Nuc | ID 3D seg. (EM nucleus) |
| Volumetric denoising (Chen et al.) | 187 | 10.1101/2025.08.26.672334 | FIB-SEM (?) | Mitochondrion | 3D | EM-Mito | – |
| Gallusser et al. | 188 | 10.1083/jcb.202208005 | FIB-SEM | Mitochondrion | 3D | EM-Mito | ID 3D seg. (mito) |
| 3D cell cultures (D'Imprima et al.) | 189 | 10.1016/j.devcel.2023.03.001 | FIB-SEM | Mitochondrion, nucleus | 3D | EM-Mito; EM-Nuc | ID 3D seg. (EM nucleus) |
| vEM data reuse (Czymmek et al.) | 190 | 10.1038/s41556-024-01381-3 | vEM (?) | Mitochondrion | 3D | EM-Mito | – |
| STARD3 (Eichler et al.) | 191 | 10.1038/s44318-026-00705-3 | vEM (?) | Mitochondrion | 3D | EM-Mito | – |
| Hepatic ER / mitochondria (Parlakgül et al.) | 192 | 10.6019/EMPIAR-12017 | FIB-SEM | Mitochondrion | 3D | EM-Mito | ID 3D seg. (mito) |
| Padman et al. | 193 | 10.1083/jcb.202411138 | FIB-SEM | Mitochondrion, nucleus | 3D | EM-Mito | OOD 3D seg. (EM nucleus) |
| webKnossos mitochondria (Jiang et al.) | 194 | 10.1016/j.crmeth.2025.100989 | vEM | Mitochondrion | 3D | EM-Mito | – |
| Mouse beta cells (Müller et al.) | 195 | 10.1083/jcb.202010039 | FIB-SEM | Nucleus | 3D | EM-Nuc | – |
| Singh et al. | 196 | 10.1016/j.str.2026.07.014 | Soft X-ray tomography (?) | Nucleus | 3D | EM-Nuc | – |
| FAFB nuclei (Mu et al.) | 197 | 10.1101/2021.11.04.467197 | ssTEM | Nucleus | 3D | EM-Nuc | ID 3D seg. (EM nucleus) |
| Shaga Devan et al. | 198 | 10.1007/s00418-022-02148-3 | TEM | Nucleus | 2D | EM-Nuc | – |
| Cardiomyocyte TEM (Kiessling et al.) | 199 | 10.1016/j.crmeth.2025.101212 | TEM | Nucleus | 2D | EM-Nuc | – |
| µSAM 3D user study data | 200 | 10.5281/zenodo.14037021 | vEM (?) | Nucleus | 3D | EM-Nuc | – |
| Perez et al. | 201 | 10.3389/fnana.2014.00126 | SBF-SEM | Nucleus | 3D | EM-Nuc | – |
| NucleoNet | 202 | 10.64898/2026.04.02.713930 | Various EM | Nucleus | 2D, 3D | EM-Nuc | – |
| Szkalisity et al. | 203 | 10.1038/s44318-025-00423-2 | FIB-SEM | Mitochondrion, nucleus | 3D | EM-Nuc | OOD 3D seg. (mito) |
| BIO407 mouse cerebellum | 204 | 10.5281/zenodo.18886823 | FIB-SEM | Mitochondrion, nucleus | 3D | EM-Nuc | OOD 3D seg. (mito) |
| Human liver (Xing et al.) | 205 | 10.64898/2026.04.22.719970 | vEM | Mitochondrion, nucleus | 3D | – | OOD 3D seg. (mito, EM nucleus) |
| Neuromuscular mitochondria (Bae et al.) | 206 | 10.1002/advs.202411191 | vEM | Mitochondrion | 3D | – | OOD 3D seg. (mito) |
| Cochlear nerve mitochondria (Lu et al.) | 207 | 10.1007/s10162-024-00957-y | SBF-SEM | Mitochondrion | 3D | – | OOD 3D seg. (mito) |
| Nuclear envelope (Spiers et al.) | 208 | 10.1111/tra.12789 | SBF-SEM | Nucleus | 3D | – | OOD 3D seg. (EM nucleus) |
| Human placenta (Acharya et al.) | 209 | 10.6019/S-BIAD3408 | Electron tomography (?) | Nucleus | 3D | – | OOD 3D seg. (EM nucleus) |
| Hippocampal CA3 (Zheng et al.) | 210 | 10.1101/2025.07.09.663979 | ssEM | Nucleus | 3D | – | OOD 3D seg. (EM nucleus) |

